# Paternal obesity alters the sperm epigenome and is associated with changes in the placental transcriptome and cellular composition

**DOI:** 10.1101/2022.08.30.503982

**Authors:** Anne-Sophie Pépin, Patrycja A. Jazwiec, Vanessa Dumeaux, Deborah M. Sloboda, Sarah Kimmins

**Affiliations:** Department of Pharmacology and Therapeutics, Faculty of Medicine, McGill University, Montreal, QC H3G 1Y6, Canada; Department of Biochemistry and Biomedical Sciences, McMaster University, Hamilton, Ontario, L8S 4K1, Canada; Departments of Anatomy & Cell Biology and Oncology, Western University, London, Ontario, N6A 3K7, Canada; Farncombe Family Digestive Health Research Institute, McMaster University Hamilton, Ontario, Canada; Departments of Obstetrics and Gynecology, and Pediatrics, McMaster University, Hamilton, Ontario, Canada; Department of Animal Science, Faculty of Agricultural and Environmental Sciences, McGill University, Montreal, QC H9X 3V9, Canada

**Keywords:** placenta, sperm, chromatin, histone, epigenetics, epigenetic inheritance, obesity

## Abstract

Paternal obesity has been implicated in adult-onset metabolic disease in offspring. However, the molecular mechanisms driving these paternal effects and the developmental processes involved remain poorly understood. One underexplored possibility is the role of paternally driven gene expression in placenta function. To address this, we investigated paternal high-fat diet-induced obesity in relation to sperm epigenetic signatures, the placenta transcriptome and cellular composition. C57BL6/J males were fed either a control or high-fat diet for 10 weeks beginning at 6 weeks of age. Males were timed-mated with control-fed C57BL6/J females to generate pregnancies, followed by collection of sperm, and placentas at embryonic day (E)14.5. Chromatin immunoprecipitation targeting histone H3 lysine 4 tri-methylation (H3K4me3) followed by sequencing (ChIP-seq) was performed on sperm to define obesity-associated changes in enrichment. Paternal obesity corresponded with altered sperm H3K4me3 enrichment at imprinted genes, and at promoters of genes involved in metabolism and development. Notably, sperm altered H3K4me3 was localized at placental enhancers and genes implicated in placental development and function. Bulk RNA-sequencing on placentas detected paternal obesity-induced sex-specific changes in gene expression associated with hypoxic processes such as angiogenesis, nutrient transport and imprinted genes. Paternal obesity was also linked to placenta development; specifically, a deconvolution analysis revealed altered trophoblast cell lineage specification. These findings implicate paternal obesity-effects on placenta development and function as one mechanism underlying offspring metabolic disease.

**Summary sentence:** Paternal obesity impacts the sperm epigenome at genes implicated in placenta development and is associated with an altered placenta transcriptome and trophoblast cell lineage specification.

## 1 Introduction

The placenta is an extraembryonic organ that regulates fetal growth and development, and contributes to long-term adult health (Regnault, Galan, Parker, & Anthony, 2002). Placental defects can result in obstetrical complications such as pre-eclampsia, stillbirth, preterm birth and fetal growth restriction (Brosens, Pijnenborg, Vercruysse, & Romero, 2011). Intrauterine growth restriction (IUGR) in turn, is associated with a heightened risk for adult-onset cardiometabolic diseases, coronary heart disease and stroke, supporting a placental role in long-term health of offspring (Brodszki, Länne, Marsál, & Ley, 2005; Crispi et al., 2010; Cruz-Lemini et al., 2016; Eriksson, Kajantie, Thornburg, & Osmond, 2016; Menendez-Castro, Rascher, & Hartner, 2018; Mierzynski et al., 2016; Morsing, Liuba, Fellman, Maršál, & Brodszki, 2014; Sarvari et al., 2017). Despite the many adverse pregnancy outcomes involving placental defects, the molecular and cellular factors that impact placental development are poorly understood (Naismith & Cox, 2021; Perez-Garcia et al., 2018). Until recently, most studies on the origins of placental pathology have focused on maternal factors. For example, placental insufficiency occurs in 10 to 15% of pregnancies, and underlying causes include advanced maternal age (Ales, Druzin, & Santini, 1990; Torous & Roberts, 2020; Wu et al., 2019), hypertension (Krielessi et al., 2012), obesity (Delhaes et al., 2018; Lutsiv, Mah, Beyene, & McDonald, 2015; MacInnis, Woolcott, McDonald, & Kuhle, 2016; Mission, Marshall, & Caughey, 2015; Sohlberg, Stephansson, Cnattingius, & Wikström, 2012), cigarette smoking (Pintican, Poienar, Strilciuc, & Mihu, 2019), drug and alcohol use, and medications (Sebastiani et al., 2018). However, emerging studies, indicate that the paternal preconception environment including diet and obesity also play a critical role in placental development and offspring health (Binder et al., 2015; Binder, Hannan, & Gardner, 2012; Jazwiec et al., 2022; Lambrot et al., 2013).

The placenta is a complex tissue arising from the differentiation of distinct cell subtypes important for its functions. In the mouse, the cells that give rise to the placenta, the trophectoderm cell lineage, first appear in the pre-implantation blastocyst at embryonic day 3.5 (E3.5). Blastocyst implantation commences at E4.5, triggering a cascade of paracrine, endocrine and immune-related events that participate in endometrial decidualization. Cells of the trophectoderm overlying the embryonic inner cell mass serve as a source of multipotent trophoblast stem cells (TSCs) that diversify as a result of spatially and epigenetically regulated transcriptional cascades, giving rise to specialized trophoblast-subtypes. The first placental fate segregation is between the extraembryonic ectoderm (EXE) and ectoplacental cone (EPC). Cells of the EPC in direct contact with the decidua give rise to the cells with invasive and endocrine capacity, including trophoblast giants (TGCs), glycocen trophoblast (GlyT), and spongiotrophoblast (SpT). Cells of the chorion will produce two layers of fused, multinucleate syncytiotrophoblast (SynT-I and SynT-II) and sinusoidal TGCs. From E8.5, the embryonic allantois becomes fused with the chorion, permitting invagination of mesoderm-derived angiogenic progenitors that form the basis of the placental vascular bed (Hemberger, Hanna, & Dean, 2020). Together, these cells form a transportive interface, the placental labyrinth zone, which is functionally critical for sustaining fetal growth throughout gestation (Rossant & Cross, 2001; Simmons & Cross, 2005). Interhemal transfer between maternal and fetal circulation commences at E10.5, and by E12.5 all terminally differentiated cell types of the mature placenta are present.

Genetic studies of placental development using mouse mutants have identified key genes for development, differentiation, maintenance and function (Perez-Garcia et al., 2018; Rossant & Cross, 2001). For example, homeobox transcription factors are required for trophoblast lineage development (e.g. *Cdx2, Eomes*) (Chawengsaksophak, James, Hammond, Köntgen, & Beck, 1997; Chen, Wang, Gong, Khoo, & Leach, 2013; Ciruna & Rossant, 1999; Kunath, Strumpf, & Rossant, 2004; Russ et al., 2000), and maintenance of SPT requires *Ascl2* and *Egfr* (Guillemot et al., 1995; Guillemot, Nagy, Auerbach, Rossant, & Joyner, 1994; Sibilia & Wagner, 1995; Tanaka, Gertsenstein, Rossant, & Nagy, 1997; Threadgill et al., 1995).

Genomic imprinting refers to monoallelic gene expression that is dependent on whether the gene was inherited maternally or paternally (Ferguson-Smith, Cattanach, Barton, Beechey, & Surani, 1991). The expression of imprinted genes is regulated by DNA methylation, acting in concert with chromatin modifications, such as histone H3 lysine 4 tri-methylation (H3K4me3) and histone H3 lysine 9 di-methylation (H3K9me2) (Dindot, Person, Strivens, Garcia, & Beaudet, 2009; McEwen & Ferguson-Smith, 2010; Wen et al., 2008). There exists 228 imprinted genes in humans and 260 in mice; many are strongly expressed in the placenta (Coan, Burton, & Ferguson-Smith, 2005; Morison, Ramsay, & Spencer, 2005; Tucci et al., 2019; Wang, Miller, Harman, Antczak, & Clark, 2013). Disruption of placental imprinting is associated with aberrant fetal growth, preeclampsia and IUGR (McMinn et al., 2006; Monk, 2015; Zadora et al., 2017). Notably, genetic manipulation studies have determined that the paternal genome is essential for extraembryonic and trophoblast development, and paternally expressed genes dominate placenta gene expression (S. C. Barton, Adams, Norris, & Surani, 1985; Sheila C. Barton, Surani, & Norris, 1984; J. McGrath & Solter, 1986; James McGrath & Solter, 1984; Surani, Barton, & Norris, 1984; Wang et al., 2013) The connection between paternal gene expression and placenta development has led to a growing interest in the role of paternal factors in placental development and function and offspring health (Wang et al., 2013). In mice, we demonstrated that paternal folate deficiency was associated with an altered sperm epigenome, differential gene expression in the placenta, and abnormal fetal development (Lambrot et al., 2013). In other mouse models, advanced paternal age and toxicant exposure have been linked to altered placental imprinting and reduced placental weight (Denomme et al., 2020; Ding, Mokshagundam, Rinaudo, Osteen, & Bruner-Tran, 2018). In human studies, recurrent pregnancy loss is associated with increased seminal reactive oxygen species (ROS) and sperm DNA damage (Jayasena et al., 2019). Male partner metabolic syndrome and being overweight have been associated with an increased risk for pre-eclampsia and negative pregnancy outcomes (Lin, Gu, & Huang, 2022; Murugappan et al., 2021). Animal models suggest that pregnancy complications that have been associated with paternal metabolic complications may be a consequence of placental dysfunction. Indeed, in mice, paternal obesity was linked to alterations in placental DNA methylation, aberrant allocation of cell lineage to trophectoderm (TE), hypoxia, abnormal vasculature, increased expression of inflammatory factors and impaired nutrient transporters (Binder et al., 2015, 2012; Jazwiec et al., 2022). These findings support the hypothesis that paternal factors impact placental development and can have negative effects on pregnancy outcomes. To explore the relationship between paternal obesity, the sperm epigenome and offspring health we previously profiled H3K4me3, a gene-activating epigenetic mark, in mouse sperm from sires fed a high-fat diet (HFD) (Pepin, Lafleur, Lambrot, Dumeaux, & Kimmins, 2022). There was an association between HFD-induced obesity, altered sperm H3K4me3, and metabolic dysfunction in offspring. However, there remains a gap in our mechanistic understanding of the connection between the sperm epigenome and offspring metabolism. Interestingly, a significant portion of genes with altered H3K4me3 in sperm after HFD were related to placental formation and function.

In the current study, we test the hypothesis that obesity-associated changes in sperm H3K4me3 drives aberrant gene expression during placental formation leading to placental dysfunction, and abnormal offspring metabolic phenotypes. To test this hypothesis, sperm was collected from obese sires and placentas from obese-sired pregnancies. Obesity-altered H3K4me3 in sperm occurred at placenta-specific enhancers and the placental transcriptome was altered in a sex-specific manner. Changes in gene expression included genes critical for placental functions that support fetal and organ system development. A deconvolution analysis revealed changes in the placental lineage specification comparable with pathological changes observed in placental defects that are associated with adult-onset cardiometabolic diseases (Aliee & Theis, 2021; Chu et al., 2019a; Cuffe et al., 2014; Han et al., 2018). Comparative analysis between placental transcriptomic profiles from our paternal HFD-induced obesity model with that of a hypoxia-induced fetal growth restriction model, revealed common placental defects across the two models (Chu et al., 2019a) consistent with our previous work that sire-obesity induces placental hypoxia (Jazwiec et al., 2022). This study revealed that paternal obesity was linked with transcriptomic and cellular defects in the placenta and may drive developmental origins of cardiometabolic disease in offspring. Confirmation of such paternal effects in humans are needed.

## 2 Results

### 2.1 High-fat diet-induced obesity alters the sperm epigenome at regions implicated in metabolism, cellular stress and placentation

Figure 1 describes the previously phenotypically characterized paternal HFD-induced obesity mouse model used in this study (Jazwiec et al., 2022). Of note, it differs from our previous model (Pepin et al., 2022) by mouse sub-strain (C57BL/6J vs C57BL/6NCrl), research setting, timing of diet exposure (at 6 vs 3 weeks of age), and the control diet (chow vs low-fat diet). This difference in experimental design allows to test the robustness of our previous results linking HFD with alteration of sperm H3K4me3. This study also newly examines functional genomic regions in relation to the placenta cell composition and transcriptomic profile including imprinted genes, placenta enhancers and transcription factor binding motifs.

**Fig. 1.**
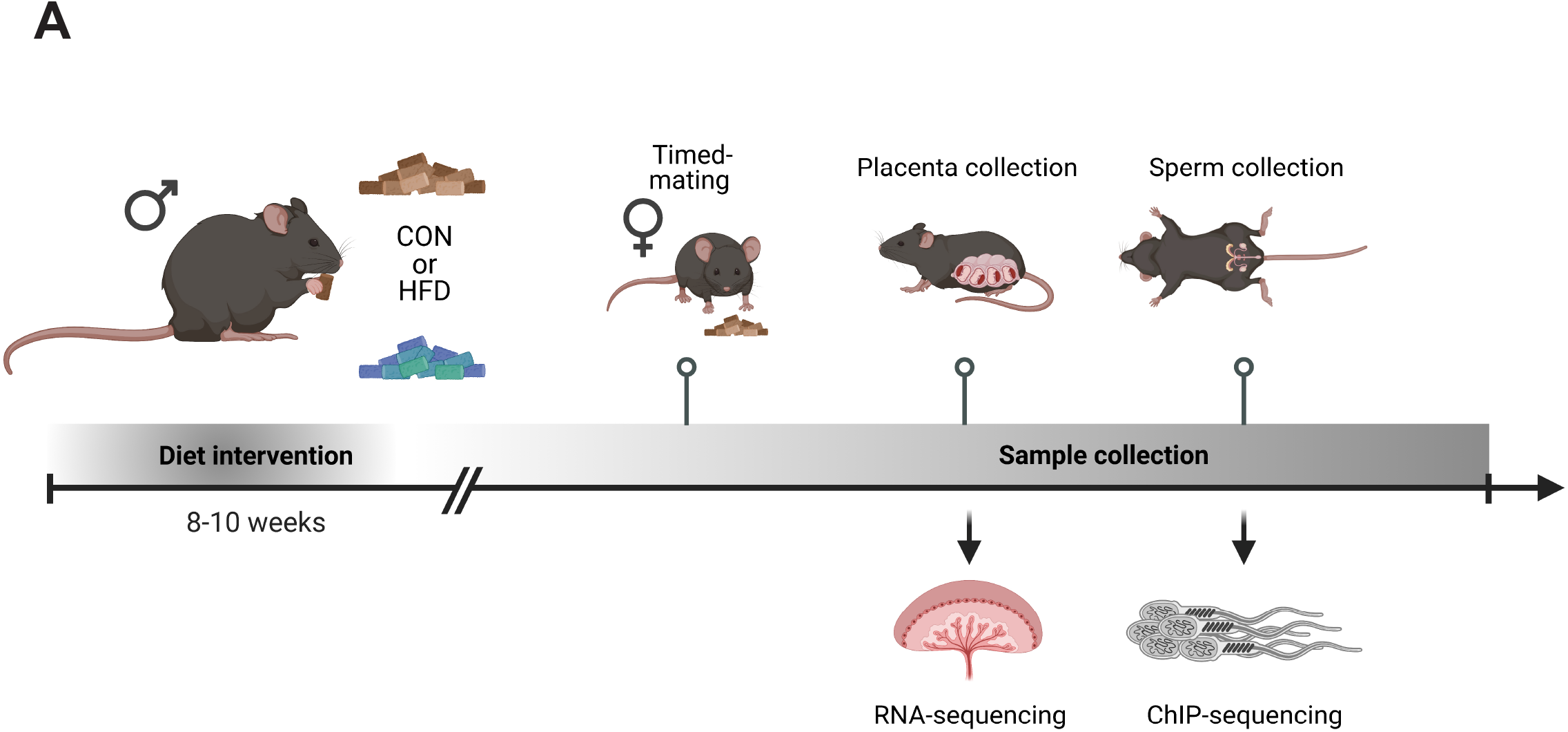
Experimental design showing the timeline and methods used to study the consequences of an obesity-induced altered sperm epigenome on the placenta. A) Six-week-old C57BL/6J sires were fed either a control or high-fat diet (CON or HFD, respectively) for 8-10 weeks. Males were then time-mated with CON-fed C57BL/6J females to generate pregnancies. Pregnant females were sacrificed at embryonic day (E)14.5 and placentas were collected to perform RNA-sequencing (RNA-seq, n=4 per sex per dietary group). Sires were sacrificed at 5 months of age and sperm from cauda epididymides was collected for chromatin immunoprecipitation sequencing (ChIP-seq, n=3 per dietary group) targeting histone H3 lysine 4 tri-methylation (H3K4me3). Created with BioRender.com.

Sperm from CON- and HFD-fed sires was profiled using ChIP-seq targeting H3K4me3 (n=3 per dietary group, Table S1). A total of 35,184 regions in sperm were enriched for H3K4me3 (Fig. S1 A; Methods), of which 28,279 were also detected in our previous study (Pepin et al., 2022). H3K4me3 profiles were highly concordant across samples which demonstrate the robustness of our profiling approach (Fig. S1 B). Principal component analysis on counts at sperm H3K4me3-enriched regions revealed separation of samples along Principal Component 1 (PC1) according to dietary treatment, after trimmed Mean of M-values (TMM) normalization and batch adjustment (Fig. S1 C). The top 5% regions (n=1,760) contributing to PC1 were considered the most sensitive to HFD-induced obesity and were selected for downstream analysis (Fig. S1 C, Fig. 2). Despite differences in experimental design and animal models, we found a significant overlap in regions showing differential H3K4me3 (deH3K4me3) from both studies (128 overlapping regions, Fisher’s exact test P=2.2e-16, Fig. S1 D). Additionally, there were similarities in terms of enriched processes between both lists of deH3K4me3 regions overlapping promoters - in particular metabolic and neurodevelopmental pathways (Fig S1 E, Table S2, Supp file 1). Consistent with our previous study, the majority of obesity-associated regions showed an increase in enrichment for H3K4me3 (n=1,257 versus n=503, 71.4%, Fig. 2 A-B). Regions losing H3K4me3 showed moderate H3K4me3-enrichment in CON sperm, with predominantly low CpG density, whereas regions gaining H3K4me3 showed low-to-moderate enrichment with mainly high CpG density (Fig. 2 C). Regions not impacted by diet showed high H3K4me3 enrichment in CON sperm, with low and high CpG density (Fig. 2 C). Consistent with our previous study, regions losing H3K4me3 were predominantly located >5 kilobase (kb) from the transcription start site (TSS), likely in intergenic spaces (Fig. 2 D i). Regions gaining H3K4me3 in HFD sperm were located near the TSS (within 1 kb), likely at promoter regions (Fig. 2 D ii). Obesity-associated deH3K4me3 regions overlapping promoters were found at genes involved in metabolic processes, cellular stress responses, vasculature development, and placentation (Fig. 2 E i-ii, Tables S3-4). Examples of genes showing deH3K4me3 in sperm include, *Cbx7* (Chromobox protein homolog 7; a component of the polycomb repressive complex 1, involved in transcriptional regulation of genes including the *Hox* gene family), *Prdx6* (Perincluded the transmembraneoxiredoxin 6; an antioxidant enzyme involved in cell redox regulation by reducing molecules such as hydrogen peroxide and fatty acid hyperoxides), and *Slc19a1* (Solute carrier family 19 member 1 or folate transporter 1; a folate organic phosphate antiporter involved in the regulation of intracellular folate concentrations) (Fig 2F). We identified deH3K4me3 in HFD-sperm at *Igf2* (Fig. 2 F ii - Insulin-like growth factor 2), a paternally-expressed imprinted gene with an essential role in promoting cellular growth and proliferation in the placenta. Importantly *Igf2* function has been related to metabolic disease and obesity (Kadakia & Josefson, 2016; Livingstone & Borai, 2014; reviewed in St-Pierre et al., 2012). Other imprinted genes with deH3K4me3 included the homeodomain-containing transcription factor *Otx2* (involved in brain and sense organs development), and the voltage-gated potassium channel *Kcnq1* gene (required for cardiac action potential).

**Fig. 2.**
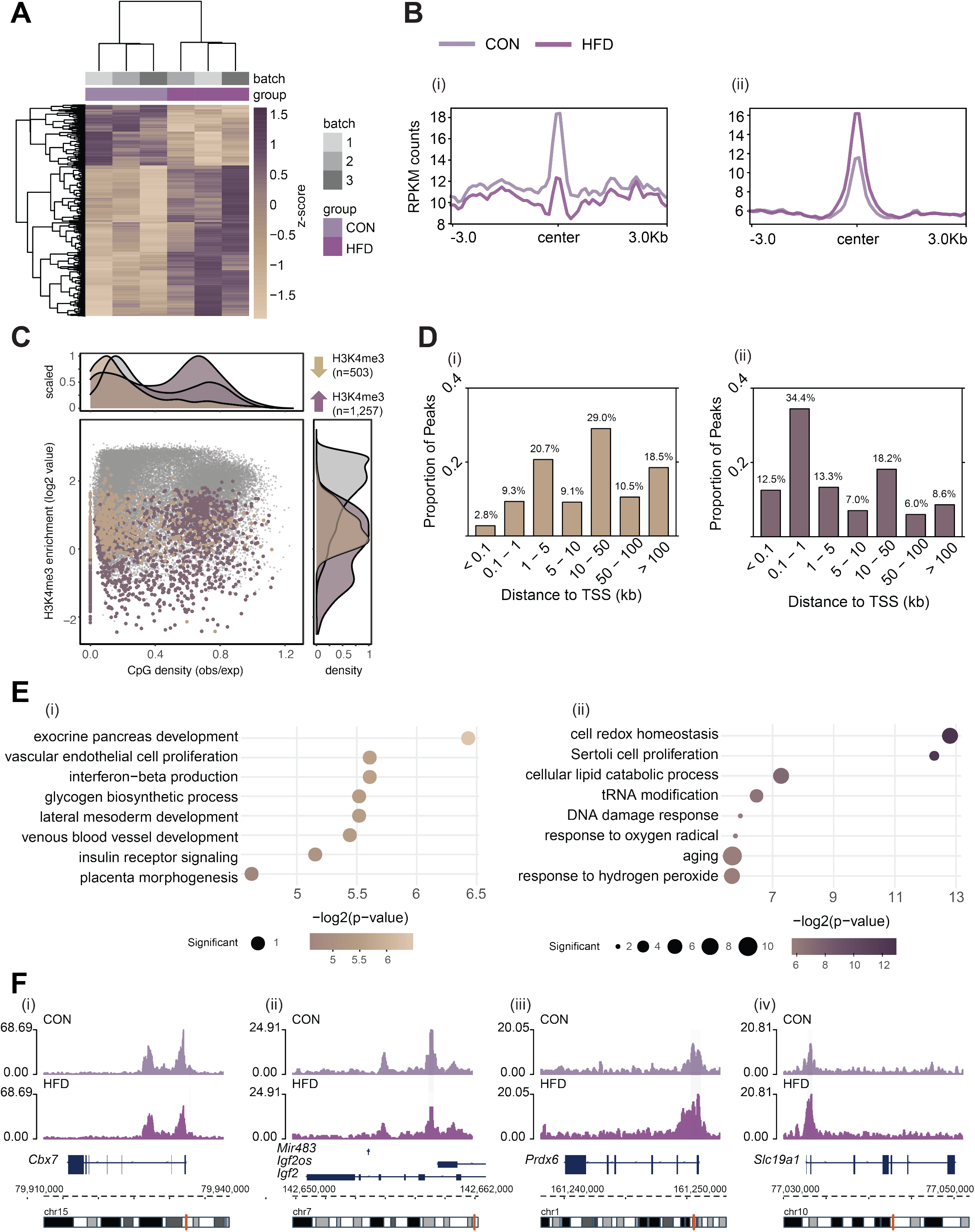
H3K4me3 signal profile at obesity-sensitive regions in sperm. A) Heatmap of log_2_ normalized counts for obesity-sensitive regions in sperm (n=1,760). Columns (samples) and rows (genomic regions) are arranged by hierarchical clustering with complete-linkage clustering based on Euclidean distance. Samples are labeled by batch (grey shades) and by dietary group. B) Profile plots showing RPKM H3K4me3 counts +/- 3 kilobase around the center of genomic regions with decreased (i) and increased (ii) H3K4me3 enrichment in HFD-sperm compared to CON-sperm. C) Scatter plot showing H3K4me3 enrichment (log_2_ counts) versus CpG density (observed/expected) for all H3K4me3-enriched regions in sperm (n=35,184, in grey), regions with HFD-induced decreased H3K4me3 enrichment (n=503, in beige), and regions with increased H3K4me3 enrichment (n=1,257, in purple). The upper and right panels represent the data points density for CpG density and H3K4me3 enrichment, respectively. D) Bar plots showing the proportion of peaks for each category of distance from the transcription start site (TSS) of the nearest gene in kilobase (kb), for obesity-sensitive regions with decreased (i) and increased (ii) H3K4me3 enrichment in HFD-sperm. E) Gene ontology (GO) analysis for promoters at obesity-sensitive regions with decreased (i) and increased (ii) H3K4me3 enrichment in HFD-sperm. The bubble plot highlights 8 significantly enriched GO terms, with their −log_2_(p-value) depicted on the y-axis and with the color gradient. The size of the bubbles represents the number of significant genes annotated to a specific GO term. Tables S3-4 include the full lists of significant GO terms. F) Genome browser snapshots showing genes with altered sperm H3K4me3 at promoter regions (CON pale purple, HFD dark purple).

### 2.2 Differentially enriched H3K4me3 in HFD sperm occurred at enhancers involved in placenta development, and at transcription factor binding sites

We previously showed that changes in sperm H3K4me3 associate with altered embryonic gene expression (Lismer, Dumeaux, et al., 2021). To gain functional insight into how deH3K4me3 in sperm may impact embryonic gene expression, we assessed the association between deH3K4me3 and tissue-specific and embryonic enhancers. Notably, deH3K4me3 localized at enhancers implicated in gene regulation of the testes, placenta, and embryonic stem cells (Fig. S2 A-B) (Shen et al., 2012). Interestingly, when searching for closest genes potentially regulated by placenta-specific enhancers, 3 were paternally-expressed imprinted genes (Tucci et al., 2019). These included the transmembrane protein *Tmem174,* the zinc finger protein *Plagl1* (a suppressor of cell growth), and the growth factor *Pdgfb* (a member of the protein family of platelet-derived and vascular endothelial growth factors; plays essential roles in embryonic development, cellular proliferation and migration).

Since H3K4me3 often localizes to promoters and can serve in the recruitment of transcription factors (TFs) (Cano-Rodriguez et al., 2016; Lauberth et al., 2013), we asked whether deH3K4me3 were significantly enriched in known TF binding site locations across the genome. Changes in H3K4me3 at these specific locations in sperm could impact embryonic gene expression – for example TFs, such as *Foxa1*, maintain an open chromatin state from the sperm to the embryo on the paternal chromatin (Jung et al., 2019, 2017). To explore this possibility, we searched for known TF binding motifs enriched in deH3K4me3 regions in sperm (Methods). The regions that gained H3K4me3 were significantly enriched for 202 TF binding motifs (P<0.05, binomial statistical test, q-value<0.05; Fig. 3 A and Supp file 2) (Heinz et al., 2010) whereas regions that had reduced H3K4me3 were not significantly enriched for TF binding motifs (q-value>0.05). Of the top 10 motifs enriched at regions with increased H3K4me3 signal in HFD-sperm, these genomic sequences were predicted to be bound by TFs belonging to the ETS, THAP, and ZF motif families (P<1e-10, q-value<0.0001; Fig. 3A). Interestingly, changes in sperm DNA methylation upon HFD feeding has been previously reported, and ETS motifs have been found to be DNA-methylation sensitive, including in spermatogonial stem cells (Domcke et al., 2015; Dura et al., 2022; Lea et al., 2018; Yin et al., 2017). Strikingly, *Sp1,* a pregnancy-specific TF associated with recurrent miscarriage, was found to be among the top TF-associated motif hits (P=1e-16, q-value<0.0001) in regions gaining H3K4me3 in sperm from obese sires (Tang et al., 2021). Furthermore, another TF of interest enriched at regions gaining H3K4me3 in HFD sperm is the Activating Transcription Factor 7 *(Atf7;* p-value=1e-3, q-value=0.0044, Supplemental File 3). Of note, this TF has been associated with oxidative stress-induced epigenetic changes in male germ cells in a mouse model of low-protein diet (Yoshida et al., 2020).

**Fig. 3.**
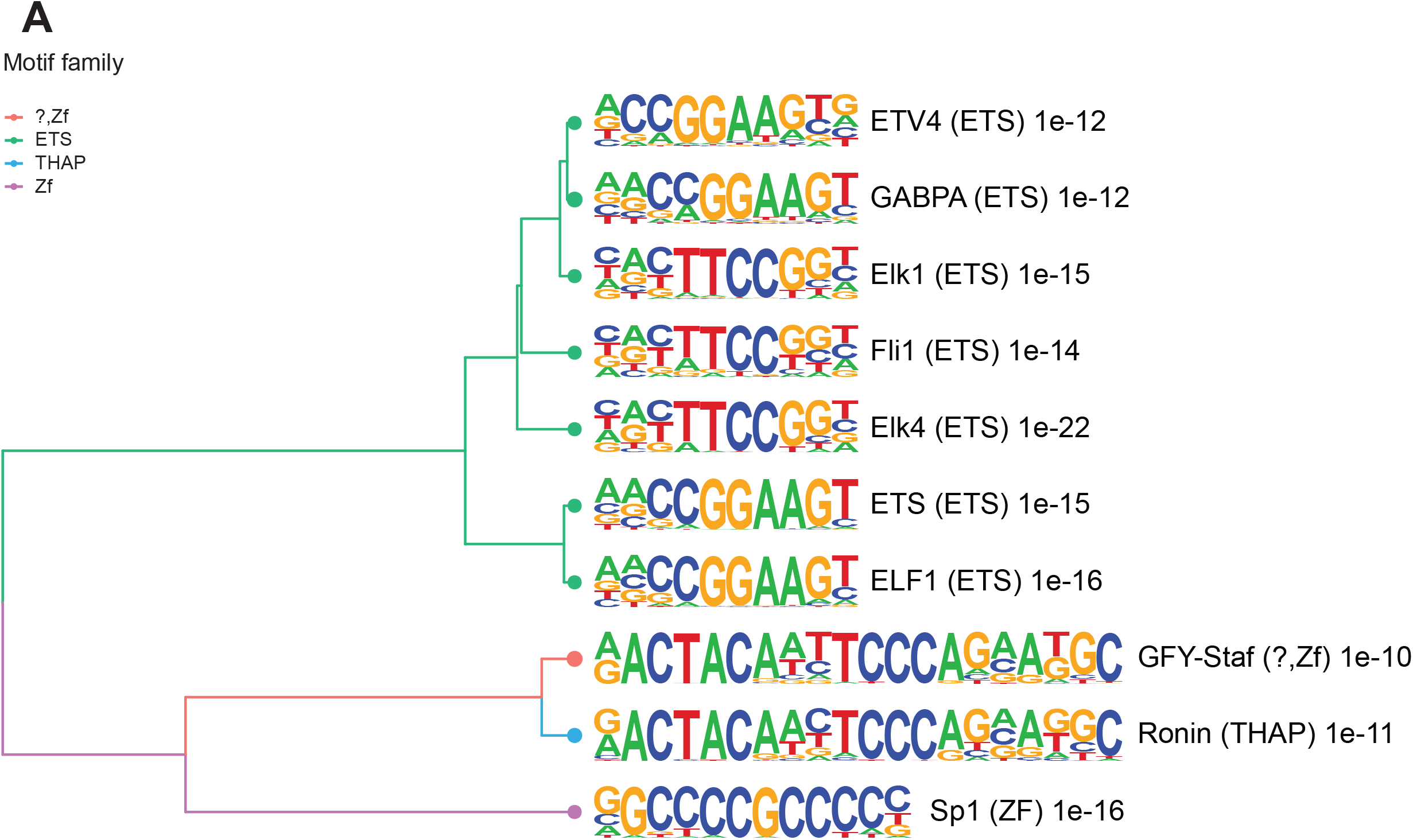
Enriched motifs at obesity-sensitive regions in sperm. A) Top 10 significantly enriched known motifs at obesity-sensitive regions with increased H3K4me3 enrichment in HFD-sperm. Motifs are clustered based on sequence similarity with hierarchical clustering. Branches of the dendrogram tree are color-coded by motif family. The name of the motif is indicated on the right, with the motif family in parenthesis, and the associated p-value for enrichment significance (binomial statistical test). The full list of enriched motifs can be found in Supplemental files 2.

Taken together we have shown there is consistency in the impacts of HFD on sperm H3K4me3 and in this model we extended our findings with a deeper functional analysis. Namely we identified novel functional genomic regions including enhancers, imprinted genes and transcription factor binding sites with altered H3K4me3 that are likely connected to paternal transmission of metabolic disease in offspring.

### 2.3 Placental gene expression is altered by paternal high-fat diet-induced obesity in a sex-specific manner

As deH3K4me3 in sperm was located at genes involved in placental formation (Fig. 2 E and Pepin et al., 2022), we assessed whether paternal obesity was associated with changes in gene expression of the placenta. We isolated RNA from E14.5 placentas derived from CON- or HFD-fed sires and performed RNA-sequencing (RNA-seq), yielding high quality data (Spearman correlation coefficient >0.89; Fig. S3 A-C). In response to paternal obesity, we detected 2,035 and 2,365 differentially expressed genes (DEGs) in female and male placentas, respectively (Fig. 4 A-B). These dysregulated genes were significantly enriched in pathways related to placental function, such as cholesterol, vitamin and protein transport, transcriptional and mRNA splicing processes, angiogenesis, and organ growth (Fig. 4 C-D, Tables S5-6). Perhaps reflecting the brain-placenta axis (Hemberger, Hanna, & Dean, 2020; Rosenfeld, 2021), other significantly enriched processes were implicated in brain and neuron development (Rosenfeld, 2021). Given that correct imprinted gene expression is critical for development, particularly of the placenta, it is noteworthy that in HFD-sired placentas 23 and 28 imprinted genes were differentially expressed in female and male placentas, respectively (Fig. 4 E-F).

**Fig. 4.**
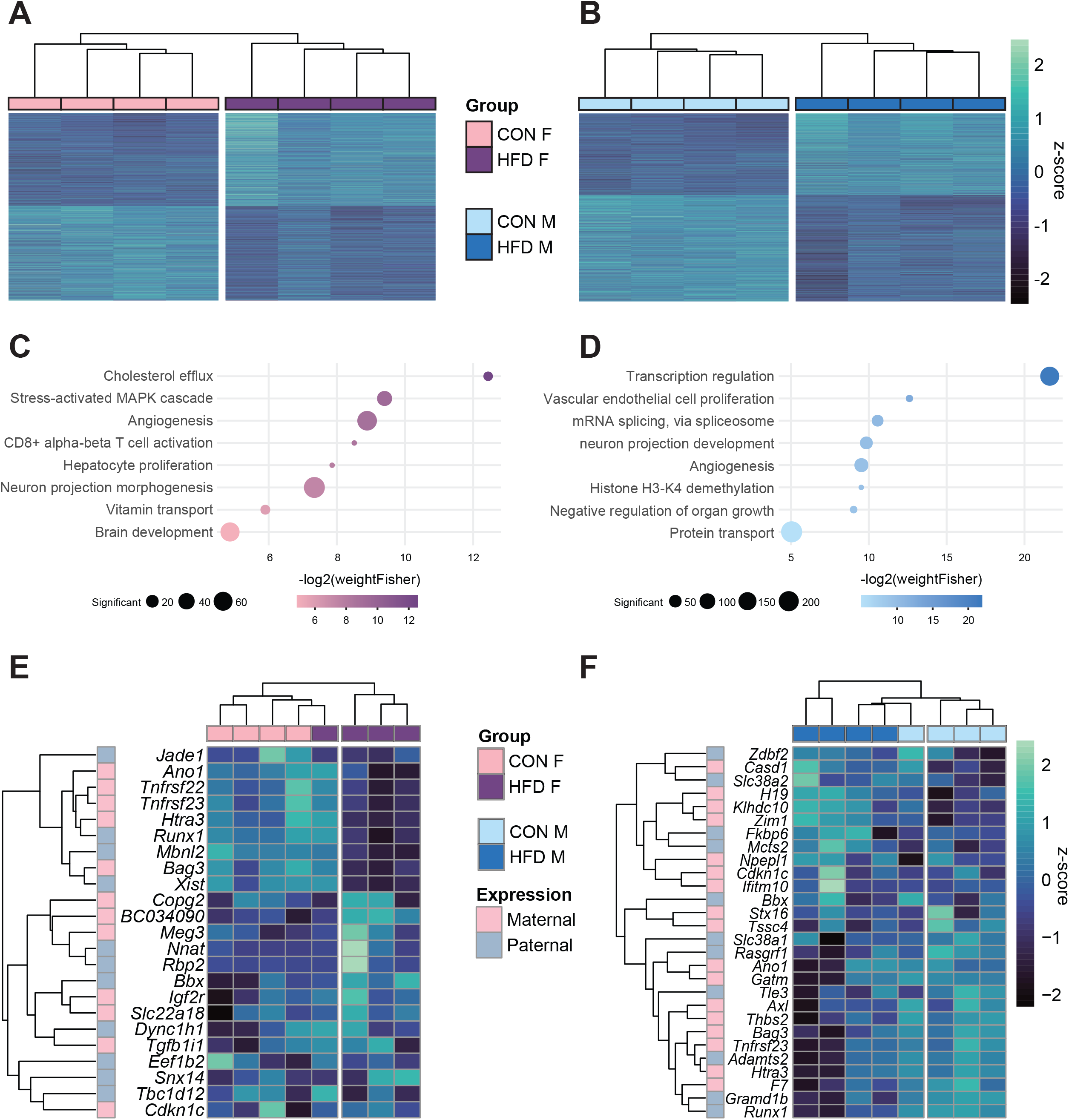
Paternal obesity alters the F_1_ placental transcriptome in a sex-specific manner. A-B) Heatmaps of normalized counts scaled by row (z-score) for transcripts that code for the detected differentially expressed genes (Lancaster p<0.05) in female (A, n=2,035 genes) and male (B, n=2,365 genes) placentas. Rows are orders by k-means clustering and columns are arranged by hierarchical clustering with complete-linkage based on Euclidean distances. C-D) Gene ontology (GO) analysis for differentially expressed genes in female (C) and male (D) placentas. The bubble plot highlights 8 significantly enriched GO terms, with their −log_2_(p-value) depicted on the y-axis and with the color gradient. The size of the bubbles represents the number of significant genes annotated to a specific enriched GO term. Tables S5-6 include the full lists of significant GO terms. E-F) Heatmaps of normalized counts scaled by row (z-score) for detected differentially expressed imprinted genes (Lancaster p<0.05) in female (E, n=23 genes) and male (F, n=28 genes) placentas. Genes are labeled based on their allelic expression (paternally expressed genes in pale grey, maternally expressed genes in pale pink). Rows are orders by k-means clustering and columns are arranged by hierarchical clustering with complete-linkage based on Euclidean distances.

Of note, although a significant number of DEGs overlapped between female and male placentas (n=359, Fisher’s exact test P=1.5e-19; Fig. S3 D i), 82% of female DEGs and 85% of male DEGs were uniquely de-regulated, indicating sex-specific placental responses to paternal obesity. The findings are concordant with previous studies which observed sex-specific effects of paternal factors on offspring metabolism (Binder et al., 2015; Claycombe-Larson, Bundy, & Roemmich, 2020; Glavas et al., 2021; Jazwiec et al., 2022). This suggests some sexually dimorphic responses may originate *in utero* due to differences in placental development and function. To assess the link between sperm H3K4me3 and the placental transcriptome, we overlapped deH3K4me3 at promoters (n=508) with DEGs in the placenta, and identified 45 and 48 DEGs in female and male placentas, respectively (Fig. S3 D ii-iii). Next, we assessed deH3K4me3 in sperm at putative placenta-specific enhancers in relation to placenta DEGs. We identified 139 putative enhancers with increased H3K4me3 and 46 with reduced H3K4me3 in HFD-sperm (Fig. S2 A-B). We then focused the analysis on the predicted genes (200 kb range) regulated by these putative enhancers (Heintzman et al., 2007; Shen et al., 2012), and defined 18 genes that were DEG in female and 19 in male placentas (Fig. S3 D iv-v). Taken together these findings show there was minor overlap between genome regulatory regions bearing deH3K4me3 and placenta DEGs. This may reflect the terminally differentiated state and heterogenous nature of the placenta at E14.5. Greater correspondence between sperm deH3K4me3 may have been observed if we had analyzed gene expression earlier in development when H3K4me3 in sperm may have a greater influence on gene expression in the first embryonic lineage of the placenta, the TE from PND 3.5. Indeed, we previously found by *in silico* analysis that there was a significant overlap between sperm and TE H3K4me3, and TE gene expression (Pepin et al., 2022). It is also worth considering that placenta profiles in this study are from bulk homogenates of whole placenta which represent a heterogeneous mixture of cell types. Bulk tissue RNA-seq measures average gene expression across these molecularly diverse cell types in distinct cellular states and the identification of DEGs can therefore be confounded by cell composition.

### 2.4 Deconvolution analysis of bulk RNA-seq reveals paternal obesity alters placental cellular composition

To assess whether there were changes in placental cellular composition associated with paternal obesity, we performed a deconvolution analysis on our bulk RNA-seq data (Fig. S4) (Aliee & Theis, 2021) using a single-cell RNA-sequencing dataset that matched the samples’ developmental stage (E14.5) and mouse strain (C57BL/6J) (Han et al., 2018). Of the 28 different cell types identified (Han et al., 2018) (Table S7; Fig. S4 A), we detected 15 cell types in our deconvolved placenta bulk RNA-seq data (Fig. 5 A and Fig. S5 A). The bulk placenta profiles were enriched for 3 trophoblast, 1 stroma and 1 endothelial cell subtypes (Figure 5A). Two of the three trophoblast cell types belonged to the spongiotrophoblast (SPT) lineage including the invasive SPT cells and SPT cells molecularly defined by highly-expressing 11-ß hydroxysteroid dehydrogenase type 2 *(Hsd11b2).* Paternal obesity was associated with changes in both SPT cell populations (Fig. 5A); we detected a significant decrease in invasive SPT cell relative abundance in female placentas (P=0.02; Fig. 5 A) and an increase in *high-Hsd11b2* SPT cells in both male and female placentas (P=0.01 and P=0.06, respectively; Fig. 5 A). These changes in SPT cellular composition indicated by this analysis upon paternal HFD-induced obesity could contribute to adult-onset metabolic dysfunction in offspring sired by obese males as observed in previous studies (Jazwiec et al., 2022; Pepin et al., 2022).

**Fig. 5.**
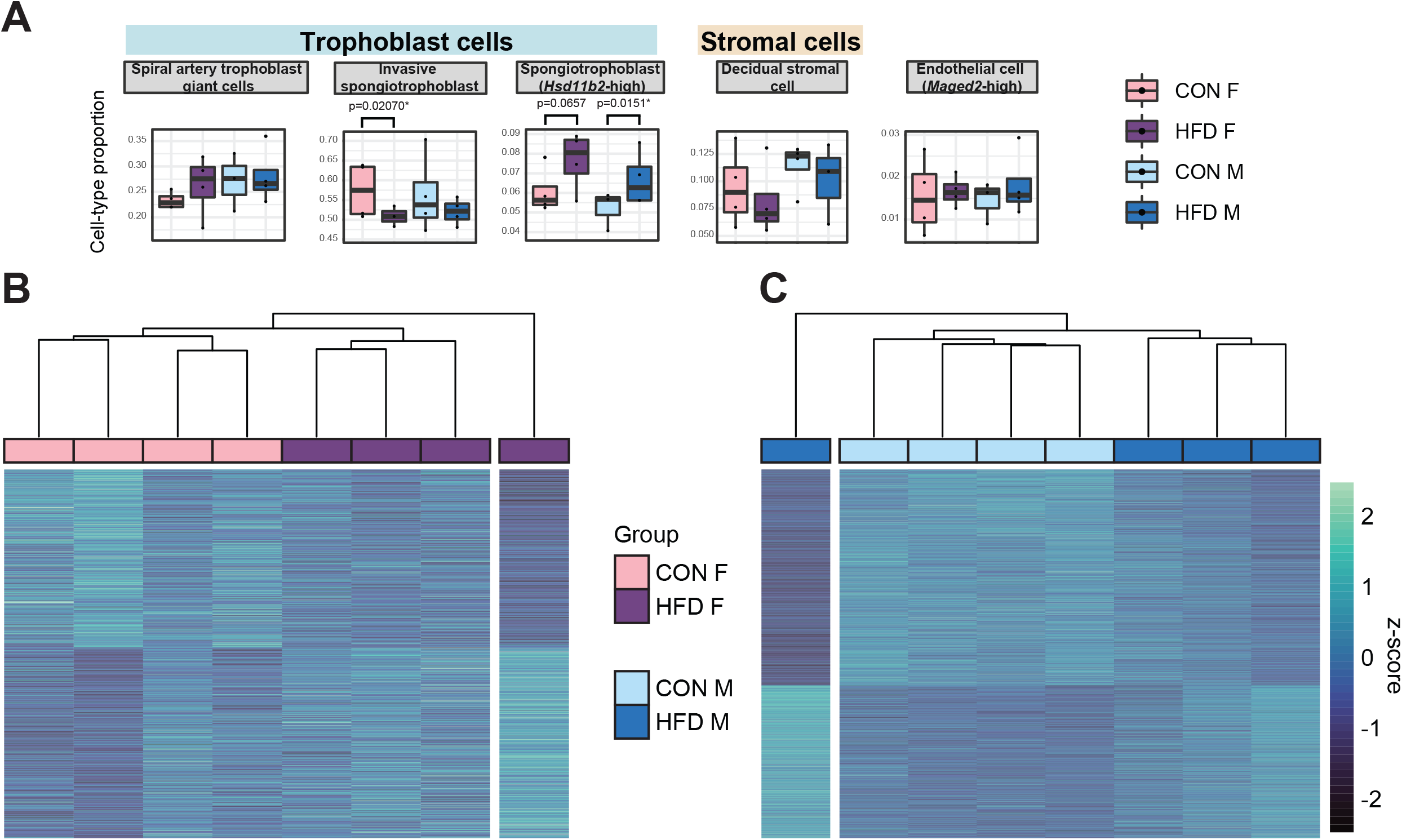
Paternal obesity-induced changes in placental cellular composition and differential expression. A) Boxplots showing sample-specific proportions for the top 5 cell types with highest proportions detected in the bulk RNA-seq data deconvolution analysis across experimental groups. Beta regression was used to assess differences in cell-type proportions associated with paternal obesity for each placental sex. P<0.05 was considered significant. B-C) Heatmaps of normalized counts scaled by row (z-score) for transcripts that code for the detected differentially expressed genes (Lancaster p<0.05) in female (B, n=423 genes) and male (C, n=1,487 genes) placentas, after adjusting for cell-type proportions. Rows are orders by k-means clustering and columns are arranged by hierarchical clustering with complete-linkage based on Euclidean distances.

To further identify gene expression changes associated with paternal obesity we performed similar differential gene expression analysis for male and female placentas but adjusted for estimated cell-type proportions (Fig. S6 A-F). We first encoded cell-type composition using the top 4 and 3 principal components identified by PCA (Fig. S6 A, B, D and E). As expected, cell types contributing the most to the sample variances for both male and female placentas included the most abundant cell types – namely invasive SPT and spiral artery TGCs, and decidual stromal cells, and endodermal cells (Fig. S6 C and F). After adjustment for placental cellular composition, we detected de-regulated genes in female (n=423 DEGS) and male placentas (n=1,487 DEGs, Fig. 5 B-C, Fig. S6 G-H), respectively. There were similarities between the bulk RNA-Seq and deconvoluted analysis in that there was overlap of DEGs detected before and after adjusting for cell-type proportions (Fig. S6 G-H). This differential gene expression analysis accounting for cellular composition provides insight into how paternal obesity may impact placental development and function.

### 2.5 Hypoxic and paternal obese-sired placentas show common transcriptomic deregulation and cell-type composition changes

Placentas derived from obese sires, like hypoxic placentas, exhibit changes in gene expression and altered angiogenesis, vasculature, and development (Binder et al., 2015, 2012; Jazwiec et al., 2022; Lin et al., 2022; McPherson et al., 2015; Mitchell et al., 2017). Hypoxia is a tightly regulated process during placental development which is essential for proper vascular formation. To determine whether paternal obese-sired placentas resemble transcriptomic and pathological phenotypes of hypoxic placentas, we compared our HFD placenta RNA-seq data to a hypoxia-induced IUGR mouse model RNA-seq data set (Chu et al., 2019a). We conducted differential gene expression analysis of the RNA-seq data from the IUGR mouse model using the same parameters as the obese-sired placenta analysis. Because this dataset did not include a sufficient number of female placenta samples, we focused the analysis on male samples only (n=5 control, n=5 hypoxic placentas). This differential analysis identified 1,935 DEGs in hypoxic placentas (Fig. S7 A-C). Likewise, we applied our deconvolution analysis described above to this bulk RNA-seq data from hypoxic placentas and detected the same principal cell types as those detected in our samples; a total of 17 different cell types were detected (Fig. 6A, Fig. S7D). Remarkably, the proportion values for each individual cell types were highly comparable across the placenta from the HFD sire model and the hypoxia mouse models (Fig. 6B). Similar to placentas derived from obese sires, hypoxic placentas showed a significant decrease in invasive SPT cell abundance (p=0.003, Fig. 6 A). Hypoxic placentas also showed a significant increase in progenitor trophoblast (*Gjb3*-high), primitive endoderm (PE) lineage (*Gkn2*-high), erythroblast (*Hbb-y*-high), and endodermal (*Afp*-high) cells, compared to control (p=0.000004, p=0.01, p=0.000003, p=0.005, respectively; Fig. 6A). Overall, the trends for directionality of changes in specific cellular abundances were consistent across the two mouse models (Fig. 6B).

**Fig. 6.**
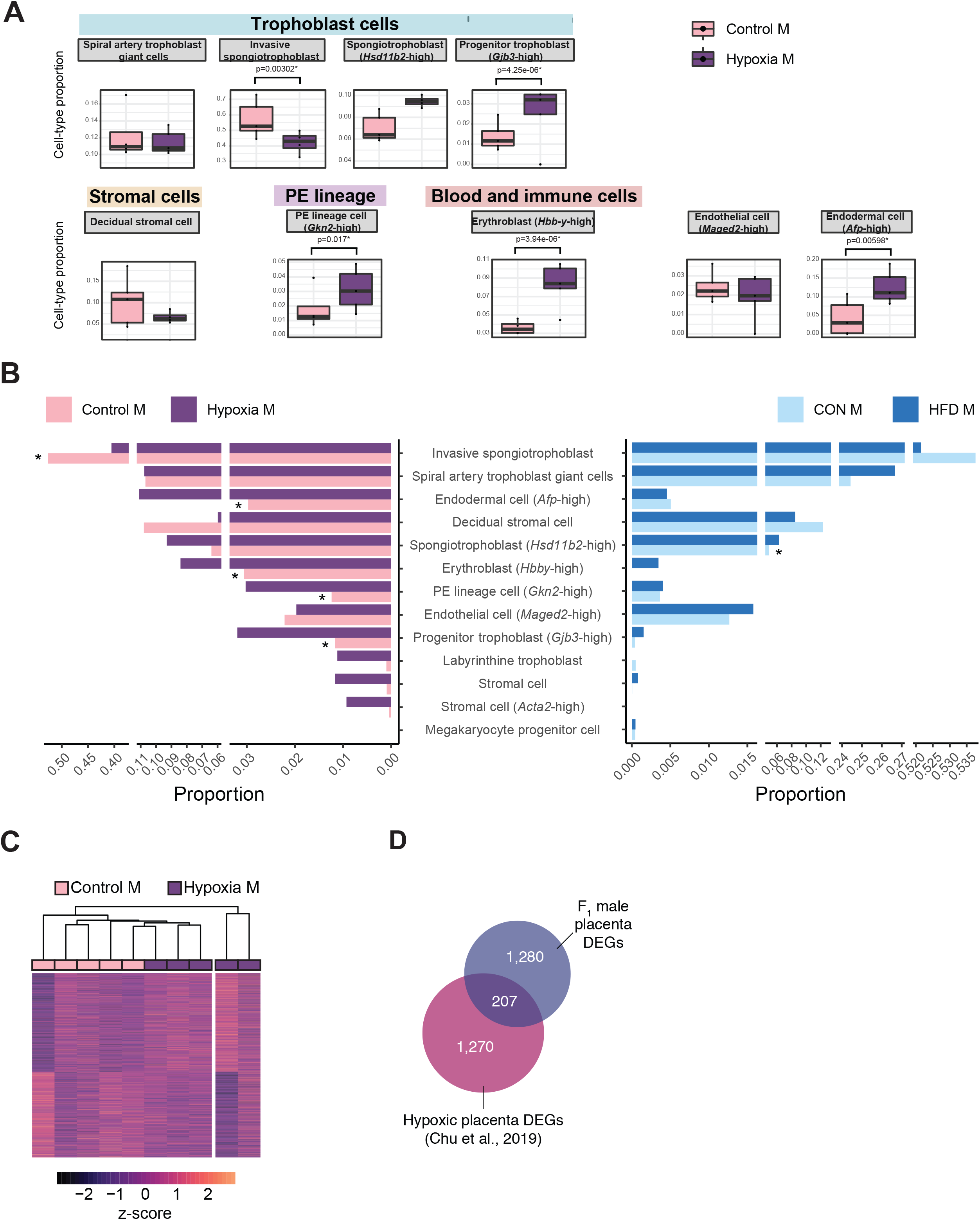
Hypoxia-induced growth restriction is associated with changes in placental cellular composition and differential expression. A) Boxplots showing sample-specific proportions for the top 10 cell types with highest proportions detected in the bulk RNA-sequencing data deconvolution analysis across experimental groups. Beta regression was used to assess differences in cell-type proportions associated with hypoxia-induced intrauterine growth restriction. P<0.05 was considered significant. B) Pyramid plot showing the median values of cell-type proportions commonly detected in both datasets assessed. The asterix (*) denote significance (P<0.05) between control versus hypoxia groups or CON M versus HFD groups, as calculated by beta regression. C) Heatmap of normalized counts scaled by row (z-score) for transcripts that code for the detected differentially expressed genes (Lancaster p<0.05, n=1,477 genes) in hypoxic placentas, after adjusting for cell-type proportions. Rows are orders by k-means clustering and columns are arranged by hierarchical clustering with complete-linkage based on Euclidean distances. D) Venn diagrams showing overlap between hypoxia-induced de-regulated genes in an intrauterine growth restriction model (Chu et al., 2019), with paternal obesity-induced de-regulated genes (this study) in male placentas.

Next, we sought to similarly investigate how much the observed changes in cellular composition within hypoxic tissues might contribute to the differential gene expression observed between conditions. Principal component analysis on placental cellular proportion values revealed a separation of samples between the control and hypoxic placentas (Fig. S7 E). Similar to the analysis of HFD-placentas, we used the top principal components (n=4 explaining 98.8% of the sample cell proportion variance) to adjust the differential expression analysis for cellular composition (Fig. S7 F). Similarly to placentas derived from HFD-fed sires, the cell types contributing the most to sample variance included the invasive SPT cells, endodermal cells (*Afp*-high), decidual stromal cells, and (Fig. S7 G). Additionally, erythroblast cells (*Hbb-y*-high) and spiral artery TGCs also strongly contributed to sample variance (Fig. S7 G). Accounting for cell-type proportions allowed for the detection of 1,477 DEGs associated with hypoxia and growth restriction (Fig. 6C), of which 356 overlapped with those initially detected before cellular composition adjustment (24%; Fig. S7 H). These data suggest that like paternal obesity-induced placental de-regulated genes, differential gene expression in hypoxic placentas is partly driven by changes in cellular composition.

Importantly, after adjusting for cell-type proportions, 207 of the paternal-obesity-induced dysregulated genes in male placentas were also found to be differentially expressed in hypoxic placentas (Fisher’s exact test P=5.1e-16; Fig. 6 D). A key gene, supporting this similarity in the molecular pathology of hypoxic placenta and obese sired placenta was the dysregulation of the imprinted gene *Igf2*. Collectively, our comparative analyses of placental transcriptomic data from both models indicate that paternal obesity, like gestational hypoxia, induces pathological and molecular consequences that are hallmarks of placental defects, and may elicit serious pregnancy complications like preeclampsia.

## 3 Discussion

Paternal health and environmental exposures impact the establishment of the sperm epigenome and are associated with altered development of the placenta, embryo, and offspring health. However, the molecular and cellular mechanisms underlying paternal obesity effects on offspring are still unclear. Our findings build on prior knowledge to show that paternal obesity alters sperm chromatin, specifically H3K4me3, in connection with widespread changes in the placental transcriptome. We provide a significant advance towards understanding the cellular and molecular drivers at the level of the sperm epigenome and placenta transcriptome that could underlie paternally-induced placental pathogenesis, growth impeded embryo development and adult-onset metabolic phenotypes.

We further observed that paternal HFD-induced obesity alters the placental transcriptome in a sex-specific manner. There is strong evidence demonstrating sex disparity in metabolic phenotypes and cardiometabolic disease risks (reviewed in Tramunt et al., 2020). These sex-specific effects are thought to be driven by sex chromosomes, hormonal factors, the gut microbiome, as well as differential fetal programming across sex in response to pre-conception and *in utero* exposures (reviewed in Sandovici, Fernandez-Twinn, Hufnagel, Constância, & Ozanne, 2022). Here, our findings suggest some of the postnatal metabolic disturbances observed in paternally-induced offspring sexually dimorphic phenotypes are established in the placenta. Interestingly, some of the de-regulated genes included imprinting genes. These genes are epigenetically controlled and inherited in a parent-of-origin manner, and the placenta is a key organ for imprinted gene function (Wood & Oakey, 2006). According to the conflict hypothesis, maternally imprinted genes (paternally expressed) support fetal growth, whereas paternally imprinted genes (maternally expressed) restrict fetal growth (Bressan et al., 2009; Haig & Graham, 1991; Moore & Haig, 1991). Some of the dysregulated imprinted genes we identified have been implicated in placental defects and pregnancy complications. For example, deletion of the gene *Htra3* (identified here as a DEG in female placentas) in mice has been implicated in IUGR owing to the disorganization of placental labyrinthine capillaries and thereby affecting offspring growth trajectories postnatally (Y. Li, Salamonsen, Hyett, Costa, & Nie, 2017). The maternally expressed gene *Copg2* (identified here as a DEG in female placentas) has been associated with pregnancies with small for gestational age infants (Kappil et al., 2015). Loss of the paternally expressed gene *Snx14* in mice (identified here as a DEG in female placentas) causes severe placental pathology involving aberrant SynT differentiation, leading to mid-gestation embryonic lethality (Bryant et al., 2020). The paternally expressed gene *Zdbf2* (DEG in male placentas) has been implicated in reduced fetal growth in mice, associated with altered appetite signals in the hypothalamic circuit (Glaser et al., 2022). Placental deficiency of the paternally expressed gene *Slc38a2* (identified here as a DEG in male placentas) leads to fetal growth restriction in mice (Vaughan et al., 2021). Lastly, mice deficient for the paternally expressed transcriptional co-repressor *Tle3* (identified here as a DEG in male placentas) show abnormal placental development including TGC differentiation failure, resulting in fetal death (Gasperowicz, Surmann-Schmitt, Hamada, Otto, & Cross, 2013). Importantly, disrupting the expression of a single imprinted gene can result in placental defect and consequently compromise fetal health or survival. It is therefore likely that the differential expression of imprinted genes detected in female and male placentas as a result of paternal obesity could at least partly explain metabolic phenotypes observed in this mouse model (Jazwiec et al., 2022; Pepin et al., 2022).

We identified changes in sperm H3K4me3 associated with paternal obesity, some of which were enriched for transcription factor binding sites. This could in turn alter TF functions. This phenomenon has been described in a mouse model of paternal low-protein diet, where oxidative stress-induced phosphorylation of the *Atf7* TF was suggested to impede its DNA-binding affinity in germ cells, leading to a decrease in H3K9me2 at target regions (Yoshida et al., 2020). As in the low-protein diet model, oxidative stress is a hallmark of obesity and increased levels of reactive oxygen species have been observed in testes of diet-induced obesity mouse models and linked to impaired embryonic development (T. Fullston et al., 2012; Lane et al., 2014; Mitchell et al., 2011). These findings provide avenues for further investigation such as whether epigenetic changes on paternal alleles may impact TF binding during early embryogenesis.

The identification of alterations of cell type proportion must be considered within the limitations of a deconvolution analysis. This analysis only provides estimates of cell-type relative within a heterogeneous tissue. This allowed us to adjust for the effect of differences in cell-type composition, but exact cell-type composition and their specific gene expression changes need to be validated by single-cell approaches such as single-cell RNA-seq or spatial transcriptomics. Furthermore, even though we used a reference dataset which included cells representative of placental tissues, the detection capacity of this approach is limited for low-abundant cell types, such as blood cells, immune cells, and inflammatory cells, which would be highly informative of placental pathological states. For example, aberrant abundance of decidual inflammatory cells, such as natural killer (NK) cells, have been linked to the pathogenesis of preeclampsia (Aneman et al., 2020; Bachmayer, Rafik Hamad, Liszka, Bremme, & Sverremark-Ekström, 2006; Du et al., 2022; Milosevic-Stevanovic et al., 2016; Williams, Bulmer, Searle, Innes, & Robson, 2009). Incidentally, it was previously shown that paternal diet-induced obesity is associated with placental inflammation (Claycombe-Larson et al., 2020; Jazwiec et al., 2022). Interestingly, many GO terms related to inflammatory processes were enriched in the obesity-induced deH3K4me3 in sperm (Fig. 2E, Table S3-4, and Pepin et al., 2022), suggesting sperm deH3K4me3 might be partly influencing placental inflammation. However due to the low representation of immune cells in the data set this could not be assessed.

### 3.1 Speculation and perspectives

Many of the DEGs in the paternal obese-sired placentas were involved in the regulation of the heart and brain. This is in line with paternal obesity associated to the developmental origins of neurological, cardiovascular, and metabolic disease in offspring (Andescavage & Limperopoulos, 2021; Binder et al., 2015, 2012; Chambers et al., 2016; Cropley et al., 2016; de Castro Barbosa et al., 2016; T. Fullston et al., 2012; Tod Fullston et al., 2013; Grandjean et al., 2015; Huypens et al., 2016; Jazwiec et al., 2022; Mitchell et al., 2011; Ng et al., 2010; Pepin et al., 2022; Perez-Garcia et al., 2018; Terashima et al., 2015; Thornburg et al., 2016; Thornburg & Marshall, 2015; Ueda et al., 2022; Wei et al., 2014). The brain-placenta and heart-placenta axes refer to their developmental linkage to the trophoblast which produces various hormones, neurotransmitters, and growth factors that are central to brain and heart development (Parrettini, Caroli, & Torlone, 2020; Rosenfeld, 2021). This is further illustrated in studies where placental pathology is linked to cardiovascular and heart abnormalities (Andescavage & Limperopoulos, 2021; Thornburg et al., 2016; Thornburg & Marshall, 2015). For example, in a study of the relationship between placental pathology and neurodevelopment of infants, possible hypoxic conditions were a significant predictor of lower Mullen Scales of Early Learning (Ueda et al., 2022). A connecting factor between the neural and cardiovascular phenotypes is the neural crest cells which make a critical contribution to the developing heart and brain (Hemberger et al., 2020; Perez-Garcia et al., 2018). Notably, neural crest cells are of ectodermal origin which arises from the TE (Prasad, Charney, & García-Castro, 2019), which is in turn governed by paternally-driven gene expression. It is worth considering the routes by which TE dysfunction may be implicated in the paternal origins of metabolic and cardiovascular disease. First, altered placenta gene expression beginning in the TE could influence the specification of neural crest cells which are a developmental adjacent cell lineage in the early embryo. TE signaling to neural crest cells could alter their downstream function. Second, altered trophoblast endocrine function will influence cardiac and neurodevelopment (Hemberger et al., 2020).

In line with these possible routes to developmental origins of obesity and metabolic disease, paternal obesity was associated with altered trophoblast lineage specification. During placentation, invasive SPT have the ability to migrate and invade the maternal-fetal interface and replace maternal vascular endothelial cells, a critical step for maternal arterial remodeling to facilitate low resistance high volume blood flow to the fetus (Silva & Serakides, 2016). Consequently, improper trophoblastic invasion has been linked to various obstetrical complications, including premature birth, fetal growth restriction, pre-eclampsia and placenta creta (Barrientos et al., 2017; Duzyj et al., 2018; O’Tierney-Ginn & Lash, 2014). Paternal obesity also induced changes in trophoblast expressing the glucocorticoid metabolizing enzyme *Hsd11b2.* In the placenta, *Hsd11b2* is responsible for the conversion of cortisol into its inactive form, cortisone, which limits fetal exposure to maternal glucocorticoid levels. Interestingly, de-regulation of *Hsd11b2* has been observed in rodent fetal growth restriction models (Chu et al., 2019a; Cuffe et al., 2014; S. et al., 2016). These aberrant cellular composition profiles suggest that paternal factors, such as diet, can induce functional changes in the placenta that mirror placental defects associated with adult-onset cardiometabolic phenotypes.

Next, it will be important to assess earlier developmental time points to determine when and how these effects originate. Indeed, studies have shown that paternal diet-induced obesity alters preimplantation development, such as cellular allocation to TE versus ICM lineages (Binder et al., 2012). Investigating multiple and earlier time points would help reveal the dynamic trajectory of paternally-induced deregulated transcriptomic and epigenetic signatures which might be at the origin of adult-onset disease. Translating these findings to humans would be beneficial to better understand the paternal preconception contribution to placental health. This is of particular relevance, given that although most obstetrical complications are thought to be rooted in the placenta, in many cases placental defects are only detected in late gestation and the etiology of these defects are oftentimes idiopathic (Hemberger et al., 2020; Regnault et al., 2002). There are no established guidelines or clinical procedures that predict pregnancy complications and placental defects associated with paternal factors. The connections we report here between paternal effects and the placental transcriptome open new avenues for the development of epigenome-based sperm diagnostics that could be used to predict pregnancy pathologies and the developmental origins of adult disease.

## 4 Methods

### 4.1 Resource availability

#### 4.1.1 Lead contact

Further information and requests for resources and reagents should be directed to and will be fulfilled by the Lead Contacts, D. Sloboda and S. Kimmins.

#### 4.1.2 Materials availability

This study did not generate new unique reagents.

#### 4.1.3 Data and code availability

The sperm H3K4me3 ChIP-Seq and placenta RNA-Seq data generated in this study are deposited at GEO under the SuperSeries GSE207326.

### 4.2 Experimental model and subject details

#### 4.2.1 Animals husbandry and dietary treatment

Animal experiments were conducted at the McMaster University Central Animal Facility, approved by the Animal Research Ethics Board, and in accordance with the Canadian Council on Animal Care guidelines. Six-week-old C57BL/6J male mice were randomly allocated to either the control (n=8; CON; standard chow diet, Harlan 8640, Teklad 22/5 Rodent Diet; 17% kcal fat, 54% kcal carbohydrates, 29% kcal protein, 3 kcal/g) or high-fat diet (n=16; HFD; Research Diets Inc., D12492; 20% kcal protein, 20% kcal carbohydrates, 60% kcal fat, 5.21 kcal/g) group, for 10-12 weeks. All animals had free access to water and food *ad libitum,* housed in the same room which was maintained at 25°C on a controlled 12-hour/12-hour light/dark cycle. After the diet intervention, male mice were housed with one or two virgin C57BL/6J females overnight. To confirm mating, females were examined the following morning, and the presence of a copulatory plug was referred to as embryonic day 0.5 (E0.5). Females confirmed as pregnant were individually housed throughout gestation and fed a standard chow diet (Harlan 8640, Teklad 22/5 Rodent Diet). Pregnant females (n=4 CON; n=5 HFD) were sacrificed at E14.5 by cervical dislocation to collect placenta samples for RNA-seq. One male and one female placenta samples per dam were collected. Placenta were cut in half, with one half snap frozen in liquid nitrogen and kept at −80°C until RNA extraction. CON-and HFD-fed male mice were sacrificed at 4-5 months of age via cervical dislocation, and sperm was collected.

### 4.3 Methods details

#### 4.3.1 Sperm isolation

Sperm was collected at necropsy from paired caudal epididymides as previously described (Hisano et al., 2013; Lismer, Lambrot, Lafleur, Dumeaux, & Kimmins, 2021; Pepin et al., 2022). Caudal epididymides were cut in 5 mL of Donners medium (25 mM NaHCO_3_, 20 mg ml^-1^ BSA, 1 mM sodium pyruvate, 0.53% vol/vol sodium DL-lactate in Donners stock), and spermatozoa were allowed to swim out by agitating the solution for 1 hour at 37°C. Sperm cells were collected by passing the solution through a 40-μm strainer (Fisher Scientific, #22363547) followed by three washes with phosphate-buffered saline (PBS). The sperm pellet was cryopreserved at −80°C in freezing medium (Irvine Scientific, cat. #90128) until used for the chromatin immunoprecipitation.

#### 4.3.2 Chromatin Immunoprecipitation, library preparation, and sequencing

Chromatin immunoprecipitation experiment was performed as previously described (Hisano et al., 2013; Lismer, Lambrot, et al., 2021; Pepin et al., 2022). In brief, samples were thawed on ice and washed with phosphate-buffered saline. Spermatozoa were counted under a microscope using a hemocytometer and 12 million cells were used per experiment. Sperm from 2-7 male mice were pooled per sample (Table S1). We used 1 M dithiothreitol (DTT, Bio Shop, cat #3483-12-3) to decondense the chromatin and N-ethylmaleimide (NEM) was used to quench the reaction. Cell lysis was performed with a lysis buffer (0.3M sucrose, 60mM KCl, 15mM Tris-HCl pH 7.5, 0.5mM DTT, 5mM McGl2, 0.1mM EGTA, 1% deoxycholate and 0.5% NP40). DNA digestion was performed in aliquots containing 2 million spermatozoa (6 aliquots per sample), with micrococcal nuclease (MNase,15 units per tube; Roche, #10107921001) in an MNase buffer (0.3 M sucrose, 85 mM Tris-HCl pH 7.5, 3mM MgCl_2_ and 2 mM CaCl_2_) for 5 minutes at 37°C. The reaction was stopped with 5 mM EDTA. Supernatants of the 6 aliquots were pooled back together for each sample after a 10 minutes centrifugation at maximum speed. A 1X solution of protease inhibitor (complete Tablets EASYpack, Roche, #04693116001) was added to each tube. Magnetic beads (DynaBeads, Protein A, Thermo Fisher Scientific, #10002D) used in subsequent steps were pre-blocked in 0.5% Bovine Serum Albumin (BSA, Sigma Aldrich, #BP1600-100) solution for 4 hours at 4°C. Pre-clearing of the chromatin was done with the pre-blocked beads for 1 hour at 4°C. Magnetic beads were allowed to bind with 5 μg of antibody (Histone H3 Lysine 4 trimethylation; H3K4me3; Cell Signaling Technology, cat. #9751) by incubating for 8 hours at 4°C. The pre-cleared chromatin was pulled down with the beads-antibody suspension overnight at 4°C. Beads-chromatin complexes were subjected to 3 rounds of washes; one wash with a low-salt buffer (50 mM Tris-HCl pH 7.5, 10 mM EDTA, 75 mM NaCl) and two washes with a high-salt buffer (50 mM Tris-HCl pH 7.5, 10 mM EDTA, 125 mM NaCl). Elution of the chromatin was done in two steps with 250 μL (2 x 125 μL) of elution buffer (0.1 M HaHCO3, 0.2% SDS, 5 mM DTT) by shaking the solution at 400 rpm for 10 minutes at 65°C, vortexing vigorously, and transferring the eluate in a clean tube. The eluate was subjected to an RNase A (5 μL, Sigma Aldrich, #10109169001) treatment shaking at 400 rpm for 1 hour at 37°C, followed by an overnight Proteinase K (5 μL, Sigma Aldrich, #P2308) treatment at 55°C. The *ChIP DNA Clean and Concentrator* (Zymo Research, #D5201) kit was used following the manufacturer’s protocol to purify the eluted DNA with 25 μL of the provided elution buffer. Libraries were prepared and sequenced at the McGill University and *Génome Québec* Innovative Centre, with single-end 100 base-pair reads on the illumina HiSeq 2500 sequencing platform (n=3 pooled samples per diet group, Table S1).

#### 4.3.3 RNA extraction, library preparation and sequencing

Extraction of RNA from placentas was performed using the RNeasy Mini Kit (Qiagen, cat. #74104) following the manufacturer’s protocol. In brief, 10-20 mg of frozen placenta were cut on dry ice. Samples were lysed in a denaturing buffer and homogenized with homogenizer pestles. Lysates were centrifuged, supernatants transferred into a clean tube, and 70% ethanol was added to lysates. An additional DNase digestion step was performed to avoid DNA contamination. Spin columns were washed twice, and total RNA was eluted with 30 μL of RNase-free water. Libraries were prepared and sequenced at the McGill Genome Centre with paired-end 100 base-pair reads on the illumina NovaSeq 6000 sequencing platform (n=4 per sex per diet group).

#### 4.3.4 Pre-processing

##### 4.3.4.1 Sperm ChIP-Sequencing data

Pre-processing of the data was performed as previously described (Pepin et al., 2022). Sequencing reads were trimmed using the *Trimmomatic* package (version 0.36) on single-end mode filtering out adapters and low-quality reads (parameters: ILLUMINACLIP:2:30:15 LEADING:30 TRAILING:30) (Bolger, Lohse, & Usadel, 2014). Reads were aligned to the mouse genome assembly *(Mus Musculus,* mm10) with *Bowtie2* (version 2.3.4) (Salzberg, 2013). *SAMtools* (version 1.9) was used to filter out unmapped reads and *Perlcode* to remove reads with more than 3 mismatches (H. Li et al., 2009). BAM coverage files (BigWig) files were created with *deeptools2* (version 3.2.1) (parameters: -of bigwig -bs 25 -p 20 --normalizeUsing RPKM -e 160 --ignoreForNormalization chrX) (Ramírez et al., 2016).

##### 4.3.4.2 Placenta RNA-Sequencing data

Sequencing data was pre-processed as previously described (Pepin et al., 2022). Sequencing reads were trimmed with *Trim Galore* (version 0.5.0) in paired-end mode to remove adapters and low-quality reads (parameters: --paired --retain_unpaired --phred33 --length 70 -q 5 --stringency 1 -e 0.1) (Krueger, 2015). Reads were aligned to the mouse reference primary assembly (GRCm38) with *hisat2* (version 2.1.0, parameters -p 8 --dta) (Kim, Langmead, & Salzberg, 2016). The generated SAM files were converted into BAM format and sorted by genomic position with *SAMtools* (version 1.9) (H. Li et al., 2009). *Stringtie* (version 2.1.2) was used to build transcripts and calculate their abundances (parameters: -p 8 -e -B -A) (Pertea et al., 2015).

##### 4.3.4.3 Publicly available datasets

Raw files for bulk RNA-sequencing in control and hypoxic placentas (n=7 and 8, respectively) were downloaded from the National Centre for Biotechnology Information (NCBI) with the Sequencing Read Archive (SRA) Toolkit (NCBI SRA: SRP137723) (Chu et al., 2019b). Files were pre-processed as described above for RNA-sequencing on single-end mode.

Processed files with raw counts for single-cell RNA-sequencing data from E14.5 mouse placenta were downloaded from NCBI (GEO: GSE108097) and metadata matrix and cluster annotations were downloaded from https://figshare.com/s/865e694ad06d5857db4b (Han et al., 2018).

### 4.4 Quantification and statistical analysis

#### 4.4.1 Visualization, statistical, and bioinformatic analyses

Bioinformatic data analyses were conducted using R (version 4.0.2) (Team, 2018) and Python (version 3.7.4) (Van Rossum, Guido; Drake, 2009). Figures were generated using the R package ggplot2 (version 3.3.3) (Wickham, 2016) and the Python package *seaborn* (version 0.9.0) (Waskom, 2021). Statistical analysis were conducted using R version 4.0.2 (Team, 2018). For all statistical tests, a p-value less than 0.05 was considered significant. To assess significance of overlap between different sets of genes, a Fisher’s exact test was performed using the *fisher.test* function from the *stats* package (version 4.0.2), and the numbers that were used to assess statistical significance were those found in the common universe (background) of both lists being compared. To assess differences in cell type proportions across experimental groups, a beta regression was performed using *betareg* function from the *betareg* package (version 3.1-4) (Ferrari & Cribari-Neto, 2004).

#### 4.4.2 Sperm ChIP-Sequencing data

ChIP-sequencing data was processed and analyzed as previously described (Pepin et al., 2022). Using *csaw* (version 1.22.1), sequencing reads were counted into 150 base-pair windows along the genome, and those with a fold-change enrichment of 4 over the number of reads in 2,000 base-pair bins were considered as genomic regions enriched with H3K4me3 in sperm (Lun & Smyth, 2016). Enriched windows less than 100 base-pair apart were merged allowing a maximum width of 5,000 base-pair (n=35,186 merged enriched regions in total). Reads were counted in those defined regions, and those with a mean count below 10 across samples were filtered out (conferring a total of n=35,184 regions). Read counts within enriched regions were normalized with TMM and corrected for batch effects arising from experimental day, using the *sva* package (version 3.36.0) (Leek, Johnson, Parker, Jaffe, & Storey, 2012; Zhang, Parmigiani, & Johnson, 2020). Spearman correlation heatmaps were generated using *corrplot* (version 0.88) and mean-average (MA) plots with *graphics* packages (Taiyun, Wei, Simko, 2021).

To detect the obesity-sensitive regions, Principal Component Analysis (PCA) was performed. We selected the top 5% regions contributing the separation of samples according to diet group along Principal Component 1 (PC1), conferring a total of 1,760 regions associated with dietary treatment. Those regions were split according to directionality change based on positive and negative log_2_ fold-change values (increased versus decreased enrichment in high-fat diet group, respectively) from the median normalized counts of each group. The selected obesity-sensitive regions were visualized with *Pheatmap* (version 1.0.12) (Kolde, 2019). Profile plots were generated using *deeptools* (Ramírez et al., 2016). The distance from the nearest transcription start site (TSS) from each selected region was calculated and visualized with *chipenrich* (version 2.12.0) (Welch et al., 2014). The genes for which their promoters overlapped the detected obesity-sensitive regions were used in the Gene Ontology (GO) analysis using *topGO* (version 2.40.0) with Biological Process ontology category and Fisher’s exact test *(weight01Fisher* algorithm) to test enrichment significance (Alexa, Rahnenführer, & Lengauer, 2006). A *weight01Fisher* p-value below 0.05 was considered significant. Genome browser snapshots of examples of detected obesity-sensitive regions were generated using *trackplot* (Pohl & Beato, 2014). Annotations for tissue-specific enhancers were downloaded from ENCODE (Shen et al., 2012) (GEO: GSE29184) and genome coordinates were converted from the mm9 to the mm10 mouse assembly using the *liftOver* function from the *rtracklayer* package (version 1.48.0) (Lawrence, Gentleman, & Carey, 2009). To determine the corresponding genes that could be regulated by tissue-specific enhancers, we scanned the landscape surrounding putative enhancer genomic coordinates, and selected the nearest gene located less than 200 kb away, given that enhancers interact with promoters located within the same domain (Heintzman et al., 2007; Shen et al., 2012). To retrieve the gene annotations, we used the function *annotateTranscripts* with the annotation database *TxDb.Mmusculus.UCSC.mm10.knownGene* (version 3.10.0) and the annotation package *org.Mm.eg.db* (version 3.11.4) from the *bumphunter* package (version 1.30.0) (Aryee et al., 2014; Jaffe et al., 2012). From the same package, the function *matchGenes* was used to annotate the putative tissue-specific enhancer genomic coordinates with the closest genes. Annotations for transposable elements and repeats were obtained from *annotatr* (version 1.14.0) (Cavalcante & Sartor, 2017) and RepeatMasker (https://www.repeatmasker.org/). Upset plots were generated using the UpSetR package (version 1.4.0) (Conway, Lex, & Gehlenborg, 2017). The motif analysis was performed using HOMER (version 4.10.4) (Heinz et al., 2010), with the binomial statistical test and standard parameters. *ViSEAGO* (version 1.2.0) (Brionne, Juanchich, & Hennequet-Antier, 2019) was used for visualization, semantic similarity and enrichment analysis of gene ontology (Fig. S1 I). Gene symbols and annotations were obtained from the *org.Mm.eg.db* database for the *Mus Musculus* species. The Biological Process ontology category was used, and statistical significance was assessed with a Fisher’s exact test with the classic algorithm. A p-value less than 0.01 was considered significant. Enriched terms are clustered by hierarchical clustering based on Wang’s semantic similarity distance and the *ward.D2* aggregation criterion.

#### 4.4.3 Placenta RNA-Sequencing data

Placenta bulk RNA-sequencing data from this study and from (Chu et al., 2019b) was processed and analyzed using the same approach, as previously described (Pepin et al., 2022). In brief, transcripts with low read counts were filtered out (mean count<10), for a total of 47,268 and 49,999 transcripts detected in male and female placentas, respectively, and 32,392 transcripts in placentas from (Chu et al., 2019b). Differential analysis was conducted with *DESeq2* (version 1.28.1) (Love, Huber, & Anders, 2014a). For the data generated in this study, we included the batch information (RNA extraction day) and dietary group in the design formula and performed a stratified analysis by running male and female samples separately (Fig. S3 B-C). For the data generated in (Chu et al., 2019b), only male samples were analyzed given there was not a sufficient number of female samples, and we included the experimental group in the formula. Independent hypothesis weighting (IHW, version 1.16) (Ignatiadis, Klaus, Zaugg, & Huber, 2016) was used for multiple testing correction and prioritization of hypothesis testing. We performed a gene-level analysis at single-transcript resolution using the Lancaster method *(aggregation* package, version 1.0.1) (Yi, Pimentel, Bray, & Pachter, 2018). This method aggregates p-values from individual transcript to detect differentially expressed genes based on changes at the transcript level. A p-value less than 0.05 was considered significant.

For visualization, variance stabilized transcript counts were used without blind dispersion estimation (Love, Huber, & Anders, 2014b). Spearman correlation heatmaps were plotted with *corrplot* (version 0.88) (Taiyun, Wei, Simko, 2021) with samples clustered by hierarchical clustering. Transcripts coding for detected differentially expressed genes were visualized with *pheatmap* (version 1.0.12) (Kolde, 2019), with samples clustered with hierarchical clustering and transcripts by k-means clustering (n kmeans=2). Gene ontology analysis was performed as described above for the sperm ChIP-seq data. For the genomic imprinting analysis, the list of known mouse imprinted genes was retrieved from (Tucci et al., 2019).

#### 4.4.4 Deconvolution analysis

We used single-cell RNA-sequencing datasets from mouse E14.5 placenta from to deconvolute our bulk RNA-sequencing data (Han et al., 2018). The following Python packages were used: *seaborn* (version 0.9.0) (Waskom, 2021), *numpy* (version 1.17.2) (Harris et al., 2020), *pandas* (version 0.25.2) (Mckinney, 2010), *pickle* (version 4.0) (Van Rossum, 2020), *scanpy* (version 1.8.2) (Wolf, Angerer, & Theis, 2018), *scipy* (version 1.7.3) (Virtanen et al., 2020), and *autogenes* (version 1.0.4) (Aliee & Theis, 2021). The *pyplot* module was loaded from the *matplotlib* library (version 3.4.2) (Hunter, 2007). The deconvolution analysis was performed following the AutoGeneS package’s available code (version 1.0.4) (Aliee & Theis, 2021). In brief, single-cell counts were log normalized and the 4,000 most highly variable genes were selected. A principal component analysis was performed (Fig. S4 A) and the cell types previously annotated in (Han et al., 2018) were visualized (Fig. S4 B). The means of each centroids for each cell type cluster was used for optimization and feature selection. AutoGeneS uses a multi-objective optimization approach to select marker genes. In this process, a search algorithm explores a set of optimal solutions (commonly called Pareto-optimal solutions) and evaluates the objective functions (in this case, correlation and distance between the cell-type specific clusters; Fig. S4 C-D). This optimization technique allows to select the 400 marker genes (Fig. S4 E). Lastly, the Nu-support vector machine (Nu-SVR) regression model (Pedregosa et al., 2011) was used to estimate the cell-type proportions for the bulk RNA-seq data from this study and from (Chu et al., 2019b). The estimated cell-type proportions were visualized as box plots for each cell type. The cell-types with percent abundance values of zero across all samples were excluded. Statistical significance across experimental groups was assessed with beta regression on the cell-types that had a median relative abundance of at least 1.5%.

#### 4.4.5 Placenta RNA-Sequencing differential analysis with cell-type proportion adjustment

To adjust for cell-type proportions in the differential analysis, while reducing the number of covariates in the model, and to account for dependence between the cell-type proportions, a principal component analysis was performed with the deconvolved cell type proportions using the *prcomp* function from R’s base statistics. The top 3 or 4 principal components were selected to capture most of the sample variance (Fig. S6 A and D, Fig. S7 E). The differential analysis described above was repeated, with the selected principal components added as covariates in the design formula to form the cell-type adjusted model.

## 5 Acknowledgements

We thank the team from Genome Quebec for the sequencing of the ChIP-seq experiment, and the team from the Applied Genomics Innovation Core of the McGill Genome Centre for the sequencing of the RNA-seq experiment.

## 6 Author Contributions

PAJ developed the murine model and provided the tissue samples. ASP performed the experiments, developed methodology, curated data, performed the formal analysis, visualization, and writing of the original draft. SK provided supervision, conceptualization, funding acquisition, and writing of the manuscript. VD advised and provided oversight of the data and statistical analysis. DMS provided supervision, conceptualization, and funding acquisition. PAJ, DMS and VD reviewed and edited the manuscript. SK is a Canada Research Chair in Epigenomics, Reproduction and Development and funding for this study is provided by the Canadian Institutes of Health Research grants to SK (DOHaD Team grant 358654 and Operating 350129) and grant (CIHR Team grant 146333; Operating 175293) to DMS. ASP is supported by scholarships from the McGill University Faculty of Medicine and Health Sciences, the Desjardins Foundation, and the McGill Centre for Research in Reproduction and Development.

## 7 Declaration of interests

The authors declare no competing interest.

## 10 Supplemental figures

**Fig. S1.**
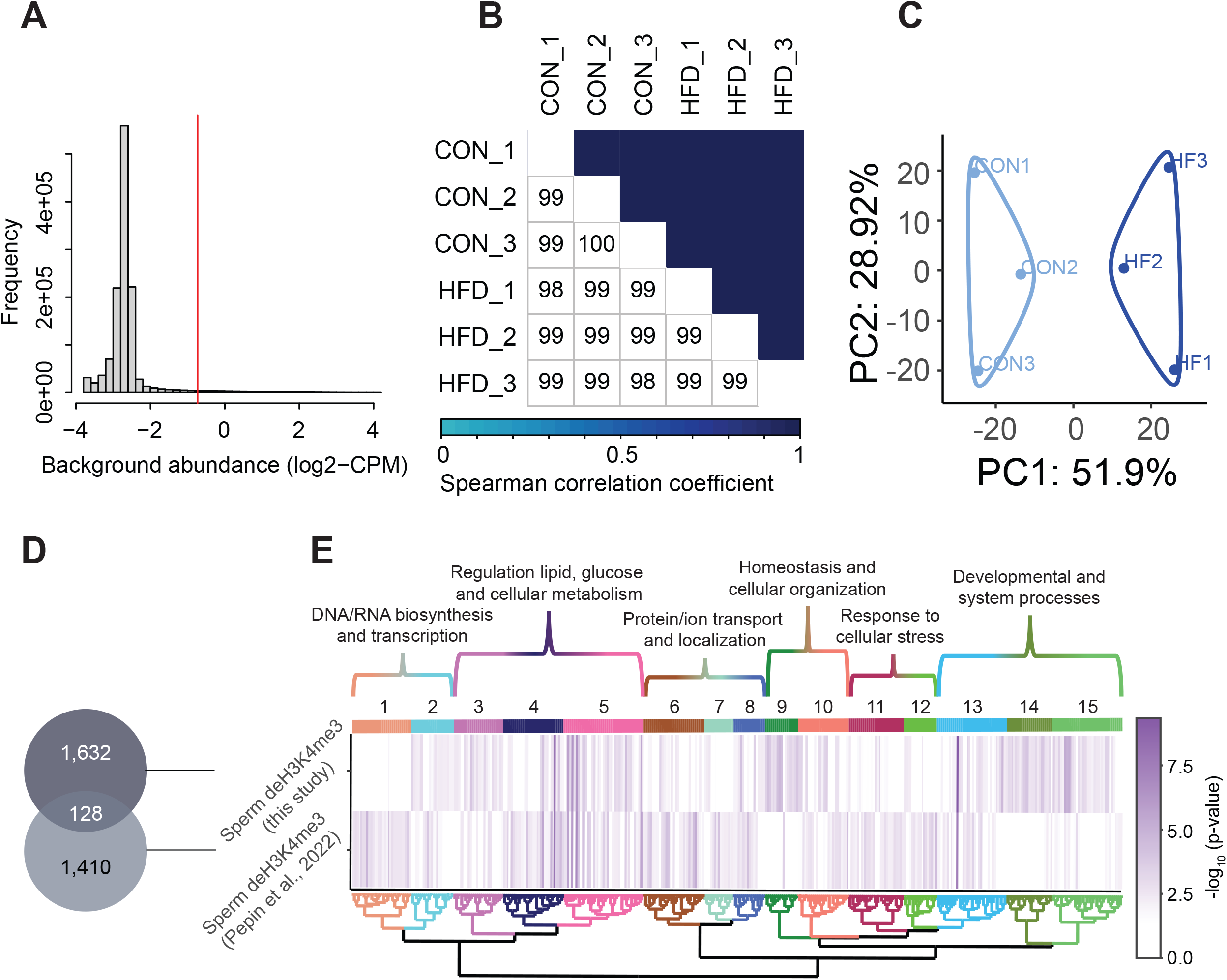
Sperm H3K4me3 ChIP-sequencing data quality and normalization. A) Histogram showing frequency distributions of read abundances of genome-wide 150 bp windows. The vertical red line indicates the cut-off where windows with low read counts were filtered out (abundance below log_2_(4) fold over 2,000 bp bins). The remaining windows (considered enriched for H3K4me3) which were less than 100 bp apart were merged allowing a maximum width of 5,000 bp (n=35,184 merged regions enriched for H3K4me3 in sperm). B) Spearman correlation heatmap on counts at sperm H3K4me3-enriched genomic regions after TMM normalization and batch adjustment. Color gradients represent correlation coefficients for each pairwise comparison. C) Principal component analysis (PCA) plot for counts in H3K4me3-enriched regions in sperm after normalization. The top 5% regions contributing to Principal Component 1 (PC1) were selected as those associated with sample separation according to dietary treatment (E). D) Venn diagram showing the overlap of detected obesity-sensitive regions from this study (dark grey) and our previous study (Pepin et al., 2022; pale grey). Significance was tested with a Fisher exact test and the p-value is shown under the graph. E) Heatmap showing significant gene ontology (GO) terms clustered based on functional similarity, comparing enriched biological functions in obesity-sensitive regions located at promoters detected in this study (top row) and in our previous study (Pepin et al., 2022, bottom row). Columns represent enriched GO terms ordered by hierarchical clustering based on Wang’s semantic similarity distance and *ward.D2* aggregation criterion. The color intensity represents the GO term enrichment significance (-log10 p-value). Interactive versions of these figures can be found in Supplemental file 1 and the complete lists of significantly enriched GO terms can be found in Table S2.

**Fig. S2.**
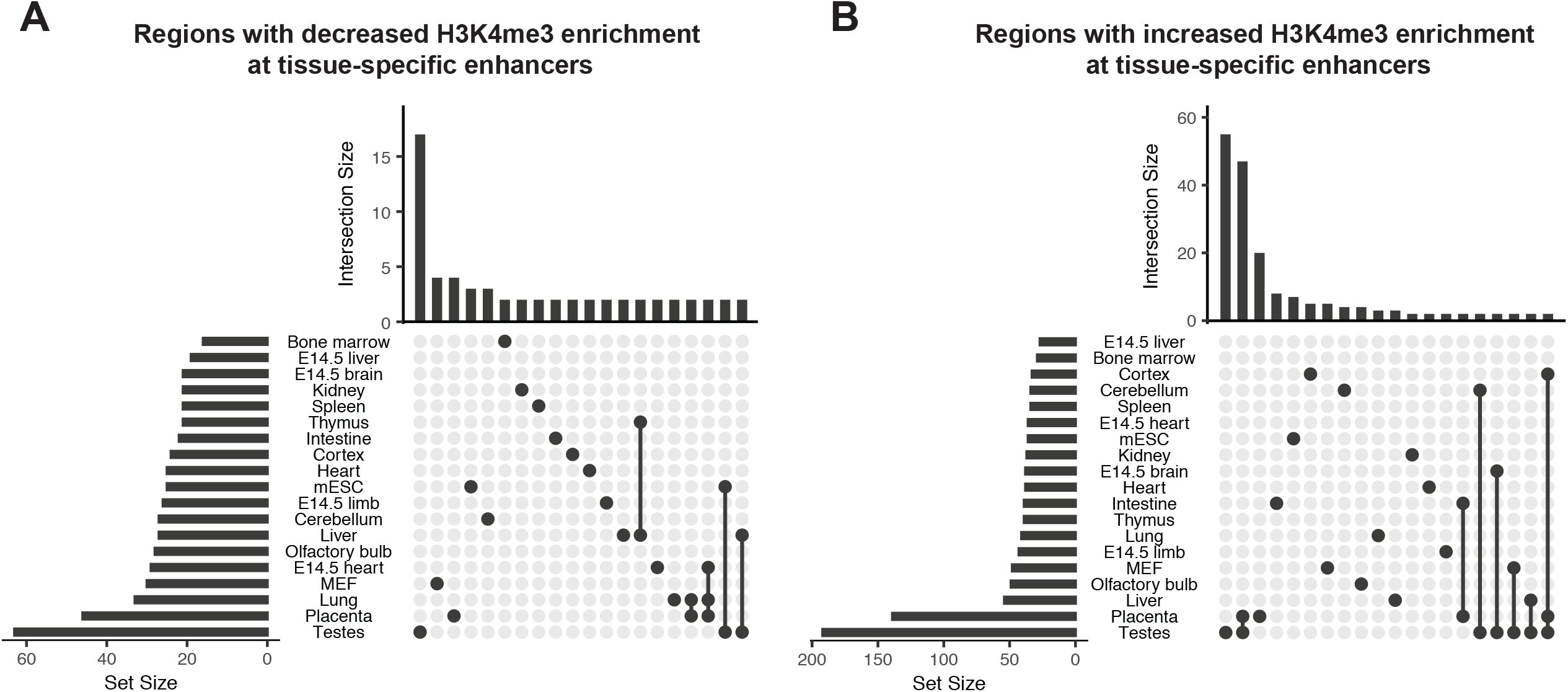
Obesity-sensitive regions in sperm are found at tissue-specific enhancers important for development. A-B) Upset plots showing annotations for tissue-specific enhancers overlapping with deH3K4me3 regions with decreased enrichment in HFD sperm (A) and increased enrichment in HFD sperm (B). Horizontal bars on the left sides of each panel represent the number of regions overlapping with each genomic annotation (set size). Vertical bars on the top of each panel represent the number of regions belonging to intersecting annotations (intersection size). Intersection sets are represented by connecting nodes.

**Fig. S3.**
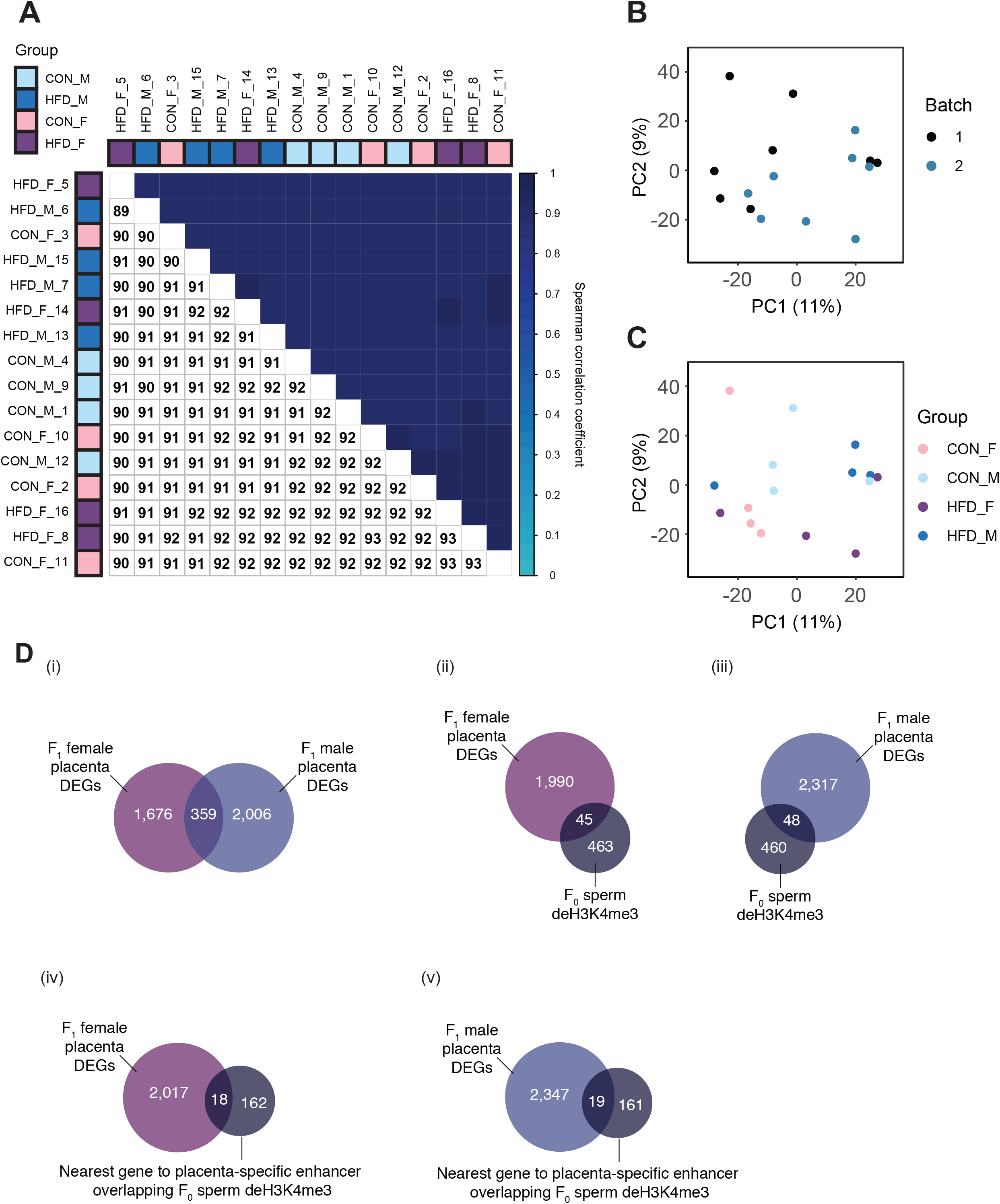
Placenta RNA-sequencing data quality assessment. A) Spearman correlation heatmap on variance stabilized transcripts. The color gradient represents the Spearman correlation coefficient for each sample pairwise comparison. B-C) Principal Component Analysis (PCA) on variance stabilized transcripts with samples labeled by batch (B) and experimental group (C). D) Venn diagrams showing the overlap of paternal obesity-induced de-regulated genes between female and male placentas (i), with sperm obesity-sensitive regions at promoters (ii and iii), and with the nearest gene to placental-specific enhancer overlapping sperm deH3K4me3 (iv and v).

**Fig. S4.**
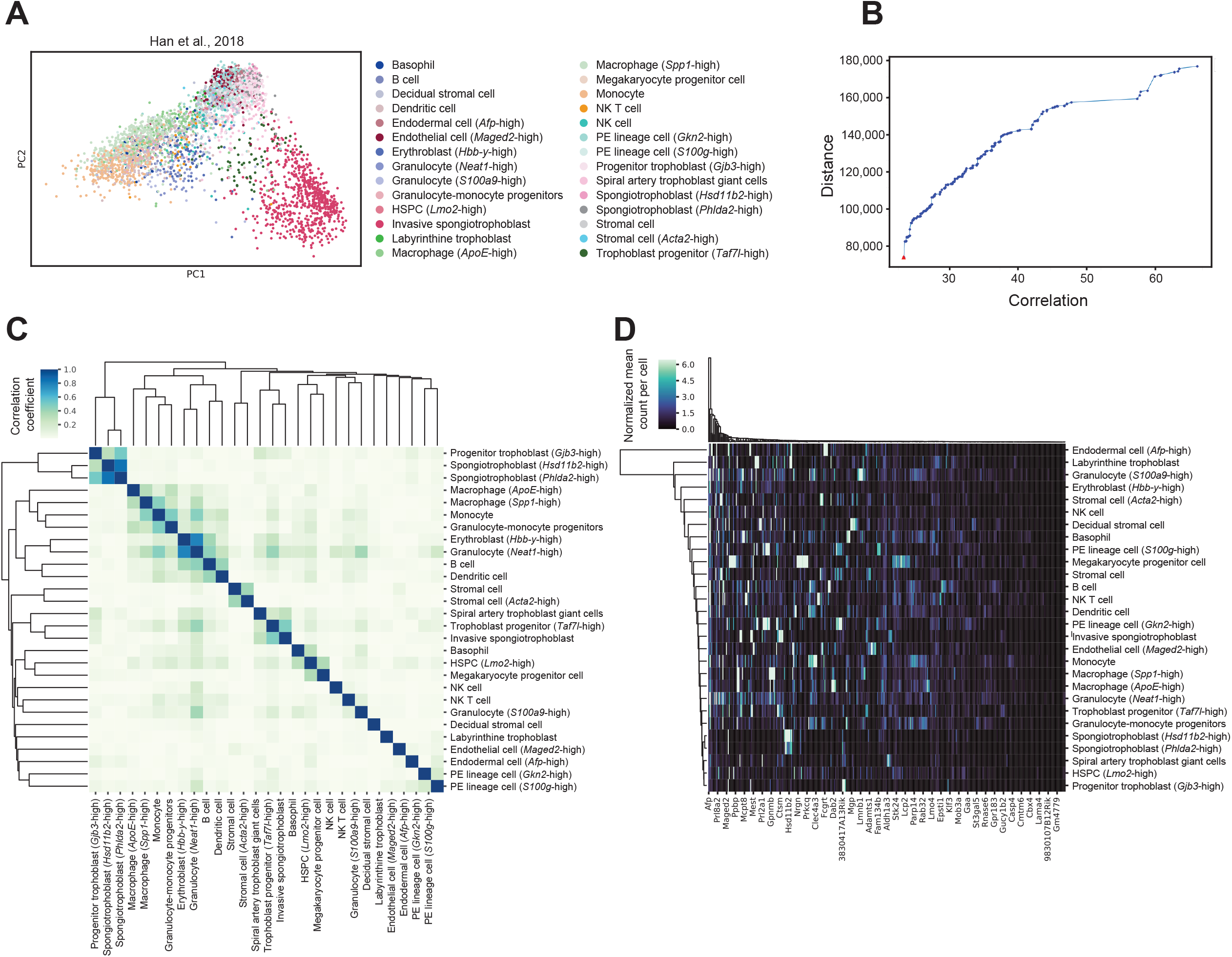
Cell-type specific marker genes selection using reference mouse E14.5 placenta single-cell RNA-sequencing dataset. A) Principal Component Analysis (PCA) plot of 4,346 single cells from mouse E14.5 placenta, with the 28 different cell types previously identified within the placenta (Han et al., 2018). The number of cells annotated to each cell type can be found in Table S7. B) The 4,000 most highly variable genes were used for feature selection using a multi-objective optimization approach with the AutoGeneS package (Aliee & Theis, 2021). The plot shows distance and correlation values for each Pareto-optimal solution. The red triangle indicates the Pareto-optimal solution used to select the 400 marker genes which maximizes distance and minimizes correlation values across cell types. C) Heatmap showing Pearson correlation between each cell-type based on expression values of the selected marker genes. The color gradient represents the Pearson correlation coefficients. Cell types are arranged by hierarchical clustering. D) Expression signatures of marker genes distinguishing the different cell types detected. The heatmap shows the mean normalized counts per cell type (rows) for the 400 marker genes (columns) as identified by AutoGeneS (Aliee & Theis, 2021). Rows and columns are arranged by hierarchical clustering.

**Fig. S5.**
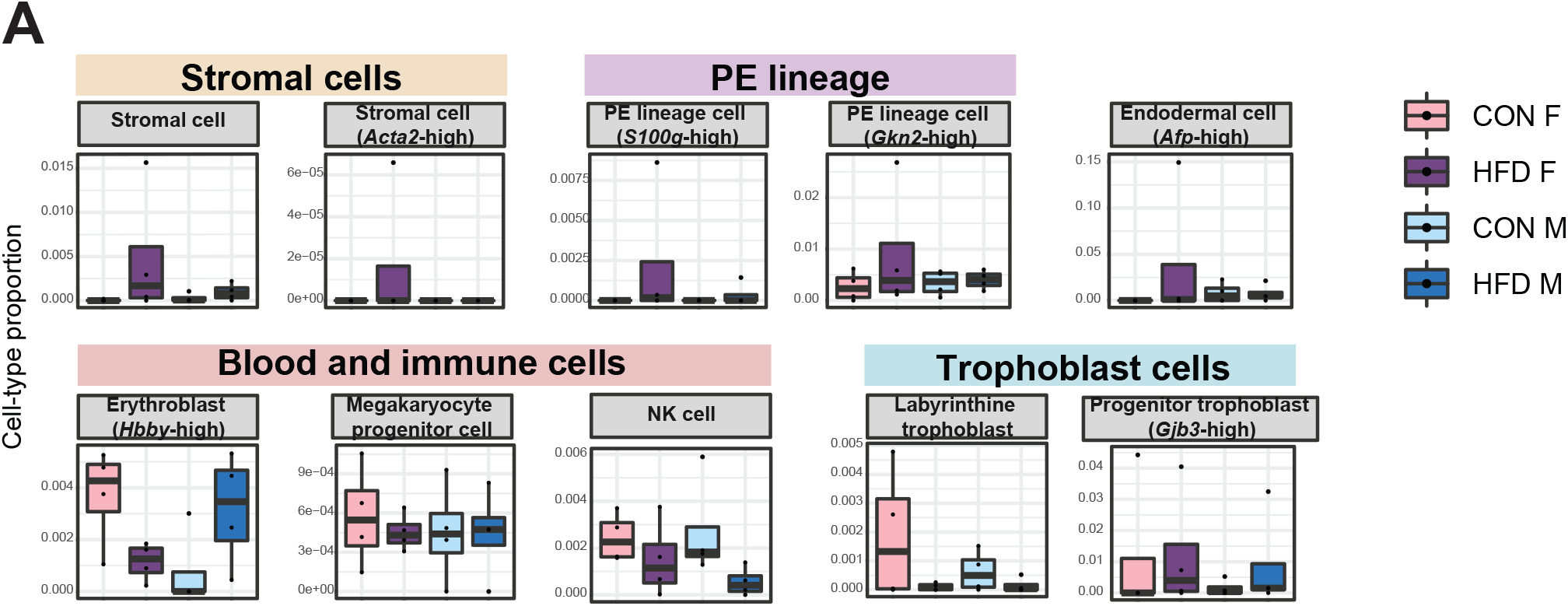
Estimated cell type proportions across experimental groups for male and female E14.5 bulk placenta tissues derived from CON- and HFD-fed sires. A) Boxplots showing sample-specific proportions for the remaining cell types detected in the bulk RNA-seq data deconvolution analysis across experimental groups. Beta regression was used to assess differences in cell-type proportions associated with paternal obesity for each placental sex. P<0.05 was considered significant.

**Fig. S6.**
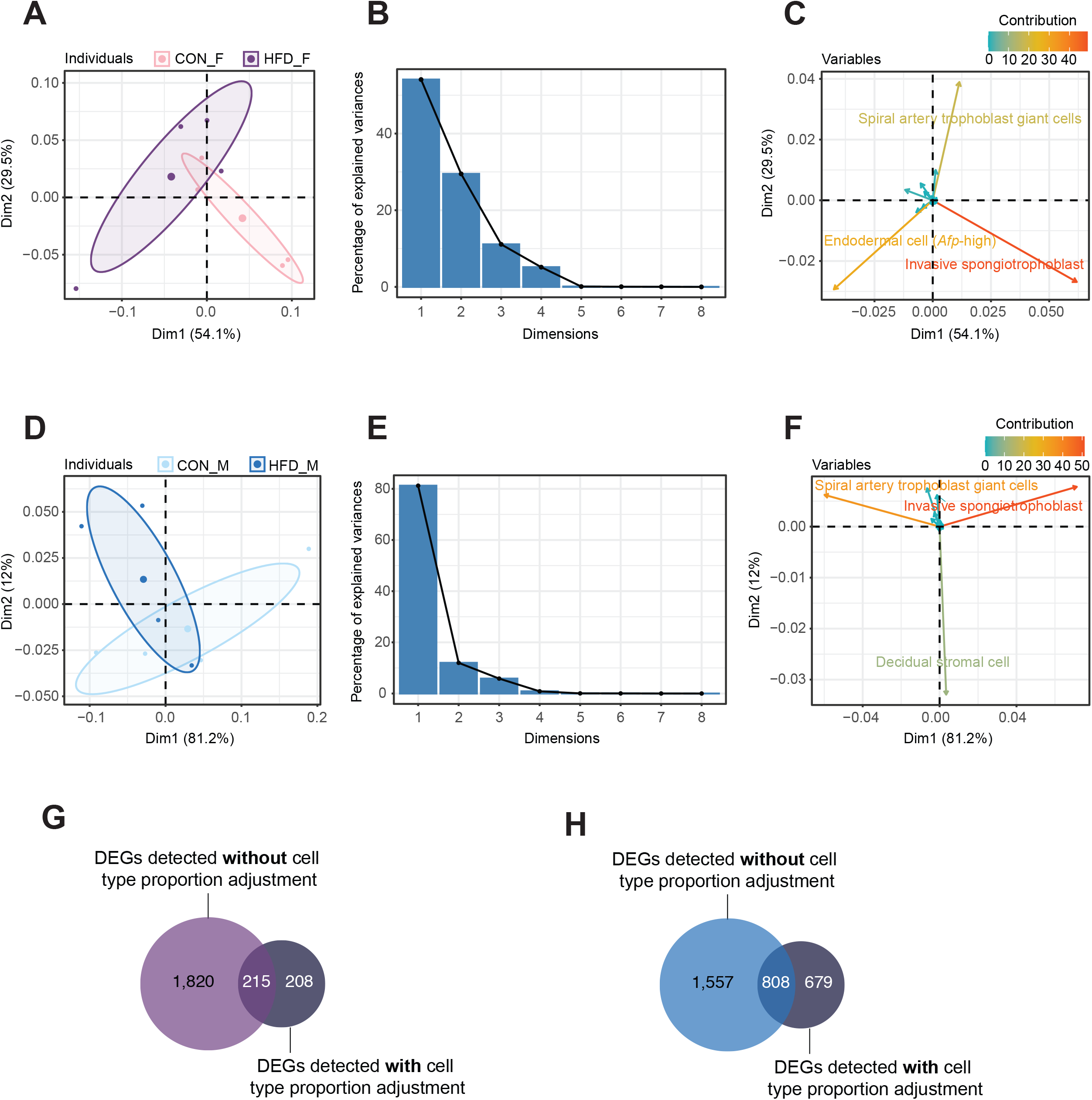
Principal component analysis (PCA) of estimated cell-type proportions. A-F) Principal component results for female (A-C) and male (D-F) placentas. A and D) Principal component analysis plot of cell-proportions. Confidence ellipses are drawn around mean points for each experimental group. B and E) Scree plots showing percentage of variances explained by each principal component (dimension). C and F) Variables factor map showing the top cell types contributing to sample variances. The color gradients on vectors represent the contribution values for each variable (cell type). G-H) Venn diagrams showing the overlap between the differentially expressed genes in female (G) and male (H) placentas, before and after adjusting for cell-type proportions.

**Fig. S7.**
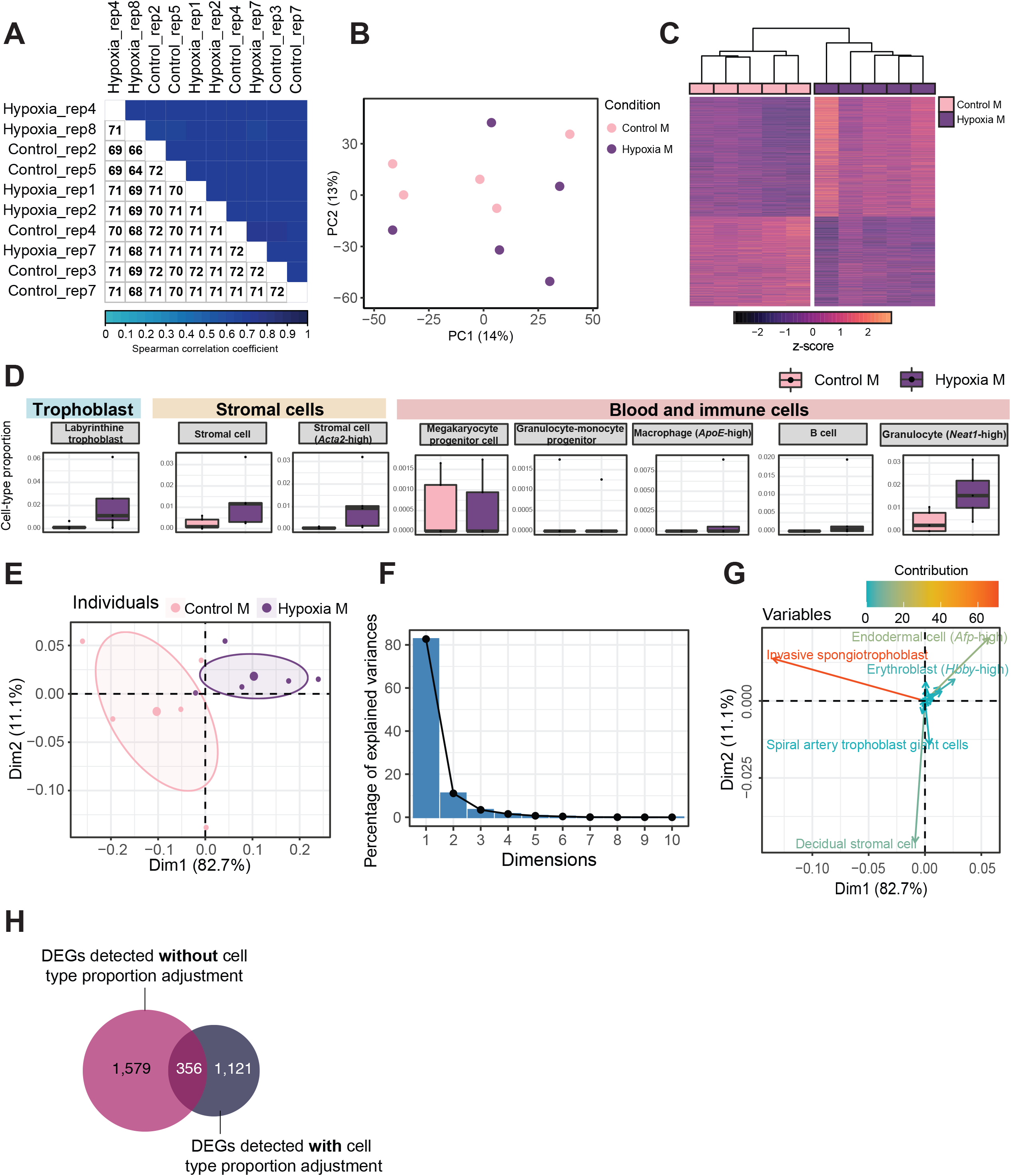
Quality assessment, processing, differential analysis, and deconvolution of RNA-sequencing data from mouse placenta in a hypoxia-induced intrauterine growth restriction mouse model. A) Spearman correlation heatmap on variance stabilized transcripts. The color gradient represents the Spearman correlation coefficient for each sample pairwise comparison. B) Principal Component Analysis (PCA) on variance stabilized transcripts with samples labeled by experimental group. C) Heatmap of normalized counts scaled by row (z-score) for transcripts that code for the detected differentially expressed genes (Lancaster p<0.05, n=1,935 genes) placentas. Rows are orders by k-means clustering and columns are arranged by hierarchical clustering with complete-linkage based on Euclidean distances. D) Boxplots showing sample-specific proportions for cell types detected in the bulk RNA-sequencing data deconvolution analysis across experimental groups. E-G) Principal component analysis of estimated cell-type proportions. E) Principal component analysis plot of cell-type proportions. Confidence ellipses are drawn around mean points for each experimental group. F) Scree plot showing percentage of variances explained by each principal component (dimension). G) Variables factor map showing the top cell types contributing to sample variances. The color gradients on vectors represent the contribution values for each variable (cell type). H) Venn diagram showing the overlap between the differentially expressed genes detected in hypoxic placentas, before and after adjusting for cell-type proportions.

## 11 Supplemental tables and files

Table S1: ChIP-sequencing sample information

Table S2: Significant gene ontology terms enriched in HFD-sperm deH3K4me3 regions at promoters detected in our previous study and this study, related to Fig S1 E

Table S3: Significant gene ontology terms enriched in HFD-sperm at regions showing a decrease in H3K4me3 at promoters, related to Fig 2 E i

Table S4: Significant gene ontology terms enriched in HFD-sperm at regions showing an increase in H3K4me3 at promoters, related to Fig 2 E ii

Table S5: Significant gene ontology terms enriched in differentially expressed genes in female placentas derived from HFD-sires, related to Fig 4 C

Table S6: Significant gene ontology terms enriched in differentially expressed genes in male placentas derived from HFD-sires, related to Fig 4 D

Table S7: Reference single-cell RNA-sequencing data information – number of cells per cell type, related to Fig S4

Supplemental file 1: Interactive heatmap for significant gene ontology terms enriched in HFD-sperm deH3K4me3 regions at promoters detected in our previous study and this study, related to Fig. S1 Ix

Supplemental file 2: Motif analysis, showing significantly enriched known motifs in regions gaining H3K4me3 in HFD-sperm, related to Fig 3 A

## Notes

### Competing Interest Statement

The authors have declared no competing interest.

