## Supplemental files 1-2 for "Paternal obesity alters the sperm epigenome and is associated with changes in the placental transcriptome and cellular composition": Supp_File_2_HOMER_up_DERs_knownResults.html

/Volumes/BINF1\_Raid/home/aspepin/projects/H3K4me3\_McMasterU/exp2/output/homer\_vs\_whole\_genome/CDvsHFD\_up/ - Homer Known Motif Enrichment Results


### Homer Known Motif Enrichment Results (/Volumes/BINF1\_Raid/home/aspepin/projects/H3K4me3\_McMasterU/exp2/output/homer\_vs\_whole\_genome/CDvsHFD\_up/)

Homer *de novo* Motif Results  
Gene Ontology Enrichment Results  
Known Motif Enrichment Results (txt file)  
Total Target Sequences = 1113, Total Background Sequences = 43621

|  |  |  |  |  |  |  |  |  |  |  |  |
| --- | --- | --- | --- | --- | --- | --- | --- | --- | --- | --- | --- |
| Rank | Motif | Name | P-value | log P-pvalue | q-value (Benjamini) | # Target Sequences with Motif | % of Targets Sequences with Motif | # Background Sequences with Motif | % of Background Sequences with Motif | Motif File | SVG |
| 1 | G A T C C T G A A G T C C G A T C G A T G A T C A G T C A C T G A T C G A G C T | Elk4(ETS)/Hela-Elk4-ChIP-Seq(GSE31477)/Homer | 1e-22 | -5.107e+01 | 0.0000 | 289.0 | 25.97% | 6394.8 | 14.66% | motif file (matrix) | svg |
| 2 | T A C G C T A G A T G C G A T C G T A C A G T C C T A G A G T C A G T C A G T C G T A C A G T C | Sp1(Zf)/Promoter/Homer | 1e-16 | -3.874e+01 | 0.0000 | 209.0 | 18.78% | 4486.7 | 10.28% | motif file (matrix) | svg |
| 3 | C T G A T G C A T A G C T G A C T A C G T C A G C T G A G C T A T C A G G A C T | ELF1(ETS)/Jurkat-ELF1-ChIP-Seq(SRA014231)/Homer | 1e-16 | -3.790e+01 | 0.0000 | 251.0 | 22.55% | 5811.4 | 13.32% | motif file (matrix) | svg |
| 4 | G A T C T C G A A G T C C G A T C G A T A G T C A T G C A C T G A T C G G A C T | Elk1(ETS)/Hela-Elk1-ChIP-Seq(GSE31477)/Homer | 1e-15 | -3.621e+01 | 0.0000 | 263.0 | 23.63% | 6275.5 | 14.39% | motif file (matrix) | svg |
| 5 | T G C A T C G A T A G C G T A C T C A G C T A G G T C A G C T A T C A G G A C T | ETS(ETS)/Promoter/Homer | 1e-15 | -3.555e+01 | 0.0000 | 175.0 | 15.72% | 3614.8 | 8.29% | motif file (matrix) | svg |
| 6 | A G T C C T G A A G T C C G A T C A G T G A T C A T G C A C T G A T C G G A C T | Fli1(ETS)/CD8-FLI-ChIP-Seq(GSE20898)/Homer | 1e-14 | -3.392e+01 | 0.0000 | 382.0 | 34.32% | 10388.7 | 23.81% | motif file (matrix) | svg |
| 7 | T C G A T C G A T A G C G T A C T C A G T A C G C G T A C G T A T C A G A G C T | GABPA(ETS)/Jurkat-GABPa-ChIP-Seq(GSE17954)/Homer | 1e-12 | -2.909e+01 | 0.0000 | 315.0 | 28.30% | 8408.1 | 19.27% | motif file (matrix) | svg |
| 8 | C T G A T A G C T G A C T C A G C T A G G T C A C G T A T C A G A G C T T C A G | ETV4(ETS)/HepG2-ETV4-ChIP-Seq(ENCODE)/Homer | 1e-12 | -2.823e+01 | 0.0000 | 388.0 | 34.86% | 11003.4 | 25.22% | motif file (matrix) | svg |
| 9 | T C A G C G T A A G T C A G C T C G T A A G T C C T G A C G T A A G T C G C A T A G T C A G T C A G T C C T G A A C T G T G C A T C G A C A T G A T C G G A T C | Ronin(THAP)/ES-Thap11-ChIP-Seq(GSE51522)/Homer | 1e-11 | -2.592e+01 | 0.0000 | 33.0 | 2.96% | 296.5 | 0.68% | motif file (matrix) | svg |
| 10 | T C G A C G T A A G T C A G C T C G T A A G T C T C G A G C T A G A C T C G A T A G T C A G T C A G T C C T G A T C A G T G C A T C G A C A G T A T C G A G T C | GFY-Staf(?,Zf)/Promoter/Homer | 1e-10 | -2.312e+01 | 0.0000 | 37.0 | 3.32% | 407.5 | 0.93% | motif file (matrix) | svg |
| 11 | T C A G T G A C G T A C T G C A G T A C C T A G G T A C A T G C A G T C G T C A A G T C G A C T | Klf9(Zf)/GBM-Klf9-ChIP-Seq(GSE62211)/Homer | 1e-9 | -2.238e+01 | 0.0000 | 197.0 | 17.70% | 4930.7 | 11.30% | motif file (matrix) | svg |
| 12 | T C A G G T A C C A T G C A G T C T G A C G T A C G A T G C A T G A C T G T C A G T A C A C T G A G T C G A C T C G A T | LBD23(LOBAS2)/colamp-LBD23-DAP-Seq(GSE60143)/Homer | 1e-9 | -2.111e+01 | 0.0000 | 298.0 | 26.77% | 8403.7 | 19.26% | motif file (matrix) | svg |
| 13 | T G C A T A G C G A C T T G C A T G A C T G C A C G T A A G C T A G C T A G T C A G T C G T A C | GFY(?)/Promoter/Homer | 1e-9 | -2.079e+01 | 0.0000 | 41.0 | 3.68% | 529.8 | 1.21% | motif file (matrix) | svg |
| 14 | G A C T T C A G C T A G A G T C A G T C G T A C A G T C C T G A A G T C A G T C A G T C G A C T A G T C A C T G A T G C | KLF3(Zf)/MEF-Klf3-ChIP-Seq(GSE44748)/Homer | 1e-9 | -2.075e+01 | 0.0000 | 235.0 | 21.11% | 6284.2 | 14.41% | motif file (matrix) | svg |
| 15 | T A C G T A C G G T A C A T C G T A C G T A C G G T C A C T G A C G T A G A C T | E2F4(E2F)/K562-E2F4-ChIP-Seq(GSE31477)/Homer | 1e-8 | -2.032e+01 | 0.0000 | 211.0 | 18.96% | 5517.6 | 12.65% | motif file (matrix) | svg |
| 16 | C T G A T C A G C A G T C T A G A C T G C T A G G A T C A T C G A C T G C T G A T C A G G A T C | Sp5(Zf)/mES-Sp5.Flag-ChIP-Seq(GSE72989)/Homer | 1e-8 | -1.995e+01 | 0.0000 | 399.0 | 35.85% | 12095.9 | 27.73% | motif file (matrix) | svg |
| 17 | G C A T G C A T G A C T G C A T T A G C A T G C G A T C C A T G A G T C A G T C | DEL2(E2FDP)/col-DEL2-DAP-Seq(GSE60143)/Homer | 1e-8 | -1.928e+01 | 0.0000 | 136.0 | 12.22% | 3185.3 | 7.30% | motif file (matrix) | svg |
| 18 | G T A C C T G A A G T C A G T C A C T G G T C A G A T C G C A T | At1g75490(AP2EREBP)/colamp-At1g75490-DAP-Seq(GSE60143)/Homer | 1e-8 | -1.927e+01 | 0.0000 | 461.0 | 41.42% | 14444.9 | 33.11% | motif file (matrix) | svg |
| 19 | T C A G T C A G G C T A C G T A T A C G G A C T T C A G T C G A C T G A C G T A T A C G G A C T | IRF8(IRF)/BMDM-IRF8-ChIP-Seq(GSE77884)/Homer | 1e-8 | -1.912e+01 | 0.0000 | 98.0 | 8.81% | 2058.6 | 4.72% | motif file (matrix) | svg |
| 20 | A T G C A G C T T C A G T G A C T C A G A T G C T G C A A C G T A T C G G A T C A C T G A G T C | NRF1(NRF)/MCF7-NRF1-ChIP-Seq(Unpublished)/Homer | 1e-8 | -1.910e+01 | 0.0000 | 97.0 | 8.72% | 2030.9 | 4.66% | motif file (matrix) | svg |
| 21 | T C G A T A G C T G C A A C T G A C T G C G T A C G T A C T A G G A C T T A C G | ETS1(ETS)/Jurkat-ETS1-ChIP-Seq(GSE17954)/Homer | 1e-8 | -1.893e+01 | 0.0000 | 315.0 | 28.30% | 9179.0 | 21.04% | motif file (matrix) | svg |
| 22 | A T G C A T G C A T C G T A C G A G C T A G T C G C T A A G T C T C A G G A C T A C T G T C G A | E-box(bHLH)/Promoter/Homer | 1e-8 | -1.847e+01 | 0.0000 | 47.0 | 4.22% | 715.9 | 1.64% | motif file (matrix) | svg |
| 23 | T C G A C T G A T A G C T G A C T C A G T C A G C G T A C G T A T C A G A G C T | ETV1(ETS)/GIST48-ETV1-ChIP-Seq(GSE22441)/Homer | 1e-8 | -1.845e+01 | 0.0000 | 393.0 | 35.31% | 12021.2 | 27.56% | motif file (matrix) | svg |
| 24 | C G A T C G A T C G A T T A C G A T C G G A T C C A T G T A G C A T G C G C T A G C T A G C T A C G T A G C A T G C A T | E2FA(E2FDP)/colamp-E2FA-DAP-Seq(GSE60143)/Homer | 1e-7 | -1.800e+01 | 0.0000 | 163.0 | 14.65% | 4107.4 | 9.42% | motif file (matrix) | svg |
| 25 | T A G C C G T A C T G A T A C G C G T A A C G T A C T G A C T G A G T C T A C G C T A G G T A C | YY1(Zf)/Promoter/Homer | 1e-7 | -1.730e+01 | 0.0000 | 50.0 | 4.49% | 818.6 | 1.88% | motif file (matrix) | svg |
| 26 | A T G C A T C G T A C G A G C T A T C G C T G A A G T C C T A G A G C T A T G C C T G A A T G C | CRE(bZIP)/Promoter/Homer | 1e-7 | -1.691e+01 | 0.0000 | 94.0 | 8.45% | 2038.4 | 4.67% | motif file (matrix) | svg |
| 27 | A T C G A G C T A C T G A G T C A C T G A G T C C G T A A C G T A C T G A G T C A C T G A G T C | NRF(NRF)/Promoter/Homer | 1e-7 | -1.667e+01 | 0.0000 | 103.0 | 9.25% | 2316.6 | 5.31% | motif file (matrix) | svg |
| 28 | C G T A C T G A G T C A A C G T A G T C C G T A A G T C A C T G A C G T A C T G T G C A A G C T | bHLH74(bHLH)/col-bHLH74-DAP-Seq(GSE60143)/Homer | 1e-7 | -1.650e+01 | 0.0000 | 92.0 | 8.27% | 1998.1 | 4.58% | motif file (matrix) | svg |
| 29 | C A T G C T A G A G T C A C T G A C T G G T A C C A T G T A C G | AT1G28160(AP2EREBP)/colamp-AT1G28160-DAP-Seq(GSE60143)/Homer | 1e-6 | -1.595e+01 | 0.0000 | 507.0 | 45.55% | 16539.8 | 37.91% | motif file (matrix) | svg |
| 30 | T G C A G T A C C T A G G A T C C A T G A C G T C G A T G C A T C G A T G C T A C G T A G C T A G T A C C T G A G A T C | CAMTA5(CAMTA)/col-CAMTA5-DAP-Seq(GSE60143)/Homer | 1e-6 | -1.593e+01 | 0.0000 | 84.0 | 7.55% | 1790.6 | 4.10% | motif file (matrix) | svg |
| 31 | C T G A A G T C C G A T A G C T A T G C G T A C A C G T A T C G C A G T G C A T | Elf4(ETS)/BMDM-Elf4-ChIP-Seq(GSE88699)/Homer | 1e-6 | -1.546e+01 | 0.0000 | 299.0 | 26.86% | 8932.6 | 20.48% | motif file (matrix) | svg |
| 32 | A T G C A T C G A T G C A T C G A T G C A T C G A T G C A T C G A T G C A T C G | SeqBias: CG-repeat | 1e-6 | -1.509e+01 | 0.0000 | 512.0 | 46.00% | 16832.4 | 38.59% | motif file (matrix) | svg |
| 33 | T G C A C T G A A G T C G T C A A C T G A C T G C G T A C G T A C T G A A G C T | EWS:FLI1-fusion(ETS)/SK\_N\_MC-EWS:FLI1-ChIP-Seq(SRA014231)/Homer | 1e-6 | -1.497e+01 | 0.0000 | 183.0 | 16.44% | 4970.4 | 11.39% | motif file (matrix) | svg |
| 34 | G C T A G C T A G C T A T C G A G T A C C T A G G A T C A C T G A C G T C A G T | CAMTA1(CAMTA)/col-CAMTA1-DAP-Seq(GSE60143)/Homer | 1e-6 | -1.465e+01 | 0.0000 | 169.0 | 15.18% | 4529.9 | 10.38% | motif file (matrix) | svg |
| 35 | C T G A A T G C C G T A A C G T A G T C A G T C A C G T A C T G A T C G G C A T | SPDEF(ETS)/VCaP-SPDEF-ChIP-Seq(SRA014231)/Homer | 1e-6 | -1.394e+01 | 0.0000 | 283.0 | 25.43% | 8514.3 | 19.52% | motif file (matrix) | svg |
| 36 | G T A C C A G T A C T G A C T G A C T G G A T C A C T G A C G T A C T G A C T G A G T C G A T C | KLF6(Zf)/PDAC-KLF6-ChIP-Seq(GSE64557)/Homer | 1e-6 | -1.394e+01 | 0.0000 | 370.0 | 33.24% | 11657.0 | 26.72% | motif file (matrix) | svg |
| 37 | A C G T C G T A T C A G A C G T A T G C C T G A T G A C C T A G C A G T T A C G G T C A A G T C | BIM3(bHLH)/col-BIM3-DAP-Seq(GSE60143)/Homer | 1e-5 | -1.346e+01 | 0.0000 | 52.0 | 4.67% | 988.1 | 2.27% | motif file (matrix) | svg |
| 38 | A C G T T A G C T G C A G A T C T C A G A C G T A C T G T G C A G A T C G A T C | Cbf1(bHLH)/Yeast-Cbf1-ChIP-Seq(GSE29506)/Homer | 1e-5 | -1.327e+01 | 0.0000 | 87.0 | 7.82% | 2005.2 | 4.60% | motif file (matrix) | svg |
| 39 | G A C T G T A C A T G C C G T A A G T C C A T G A G T C C A T G G A T C A G T C G C A T G A T C | FHY3(FAR1)/Arabidopsis-FHY3-ChIP-Seq(GSE30711)/Homer | 1e-5 | -1.266e+01 | 0.0001 | 145.0 | 13.03% | 3884.2 | 8.90% | motif file (matrix) | svg |
| 40 | A T G C A G T C C T G A A G T C C G A T A C G T A G T C A G T C A C G T A T C G G A C T A C G T | Etv2(ETS)/ES-ER71-ChIP-Seq(GSE59402)/Homer | 1e-5 | -1.249e+01 | 0.0001 | 256.0 | 23.00% | 7705.4 | 17.66% | motif file (matrix) | svg |
| 41 | T A C G T A C G G T A C A T C G A C T G T A C G T C G A C T G A T C G A A T C G | E2F6(E2F)/Hela-E2F6-ChIP-Seq(GSE31477)/Homer | 1e-5 | -1.240e+01 | 0.0001 | 215.0 | 19.32% | 6278.8 | 14.39% | motif file (matrix) | svg |
| 42 | T G A C C T G A A G T C A G T C A C T G G A T C G A C T G C A T | At5g18450(AP2EREBP)/col-At5g18450-DAP-Seq(GSE60143)/Homer | 1e-5 | -1.167e+01 | 0.0002 | 446.0 | 40.07% | 14767.6 | 33.85% | motif file (matrix) | svg |
| 43 | G A C T A T C G C T G A A G T C T C A G G A C T G T A C C T G A A G C T G T A C | TGA6(bZIP)/colamp-TGA6-DAP-Seq(GSE60143)/Homer | 1e-5 | -1.167e+01 | 0.0002 | 140.0 | 12.58% | 3794.9 | 8.70% | motif file (matrix) | svg |
| 44 | G A C T A C T G C G T A A G T C T C A G G C A T G T A C C G T A A C G T G A T C | TGA1(bZIP)/colamp-TGA1-DAP-Seq(GSE60143)/Homer | 1e-5 | -1.167e+01 | 0.0002 | 104.0 | 9.34% | 2625.3 | 6.02% | motif file (matrix) | svg |
| 45 | A G T C C T A G C T A G A G T C G A T C G T A C A G T C C T A G A G T C A G T C A G T C G T A C | Sp2(Zf)/HEK293-Sp2.eGFP-ChIP-Seq(Encode)/Homer | 1e-5 | -1.163e+01 | 0.0002 | 486.0 | 43.67% | 16289.0 | 37.34% | motif file (matrix) | svg |
| 46 | C T A G T C G A C T G A C G T A T A C G G A C T T C A G T C G A G T C A T G C A T A C G A G C T | IRF2(IRF)/Erythroblas-IRF2-ChIP-Seq(GSE36985)/Homer | 1e-4 | -1.144e+01 | 0.0002 | 36.0 | 3.23% | 630.6 | 1.45% | motif file (matrix) | svg |
| 47 | T C A G A G C T A T G C C G T A A G T C T C A G A C G T A T C G T C G A A G T C G A T C T G A C | TFE3(bHLH)/MEF-TFE3-ChIP-Seq(GSE75757)/Homer | 1e-4 | -1.131e+01 | 0.0003 | 37.0 | 3.32% | 660.9 | 1.51% | motif file (matrix) | svg |
| 48 | A C T G A C G T C A T G A T C G A T C G T G A C A C T G A T C G A T C G T G C A C T G A C G T A | E2F3(E2F)/MEF-E2F3-ChIP-Seq(GSE71376)/Homer | 1e-4 | -1.114e+01 | 0.0003 | 238.0 | 21.38% | 7210.9 | 16.53% | motif file (matrix) | svg |
| 49 | T A G C C G A T T A C G A C T G A G T C A C T G A T C G A T C G C G T A C T G A | E2F1(E2F)/Hela-E2F1-ChIP-Seq(GSE22478)/Homer | 1e-4 | -1.113e+01 | 0.0003 | 118.0 | 10.60% | 3112.2 | 7.13% | motif file (matrix) | svg |
| 50 | T C G A A C G T A C T G C T G A A G T C T C A G A G C T G T A C C G T A A G C T G A T C T C G A | JunD(bZIP)/K562-JunD-ChIP-Seq/Homer | 1e-4 | -1.111e+01 | 0.0003 | 37.0 | 3.32% | 666.3 | 1.53% | motif file (matrix) | svg |
| 51 | C G A T A C T G A G C T T G A C C T G A A G T C C T A G G C A T A T C G C G T A | SPCH(bHLH)/Seedling-SPCH-ChIP-Seq(GSE57497)/Homer | 1e-4 | -1.110e+01 | 0.0003 | 252.0 | 22.64% | 7711.4 | 17.68% | motif file (matrix) | svg |
| 52 | T C G A C T A G A G T C A G T C C G T A C G T A A C G T T A G C T C A G T A C G | NFY(CCAAT)/Promoter/Homer | 1e-4 | -1.107e+01 | 0.0003 | 193.0 | 17.34% | 5642.7 | 12.93% | motif file (matrix) | svg |
| 53 | T C G A T A G C G T C A A C T G A C T G C G T A C G T A C T A G A G C T T C A G | ERG(ETS)/VCaP-ERG-ChIP-Seq(GSE14097)/Homer | 1e-4 | -1.101e+01 | 0.0003 | 401.0 | 36.03% | 13167.3 | 30.18% | motif file (matrix) | svg |
| 54 | G A T C G C T A C A G T A C G T T A C G A G T C A T G C C T A G A G T C T C G A | Zfp57(Zf)/H1-ZFP57.HA-ChIP-Seq(GSE115387)/Homer | 1e-4 | -1.094e+01 | 0.0003 | 165.0 | 14.82% | 4694.9 | 10.76% | motif file (matrix) | svg |
| 55 | C T G A C T A G G C A T A T G C C G T A A G T C A C T G A C G T T A C G C T G A | HY5(bZIP)/colamp-HY5-DAP-Seq(GSE60143)/Homer | 1e-4 | -1.089e+01 | 0.0003 | 192.0 | 17.25% | 5625.4 | 12.90% | motif file (matrix) | svg |
| 56 | G C A T C G T A G C T A G A C T C G A T G A C T A G T C C A G T A G T C A G T C A C T G C T A G G T A C C T A G C T G A | AT5G05550(Trihelix)/col-AT5G05550-DAP-Seq(GSE60143)/Homer | 1e-4 | -1.084e+01 | 0.0003 | 383.0 | 34.41% | 12518.6 | 28.70% | motif file (matrix) | svg |
| 57 | C T G A T A C G G C A T A G C T A G C T A G T C T C G A A C T G C A G T A G C T A G C T G A T C | IRF3(IRF)/BMDM-Irf3-ChIP-Seq(GSE67343)/Homer | 1e-4 | -1.080e+01 | 0.0004 | 75.0 | 6.74% | 1772.4 | 4.06% | motif file (matrix) | svg |
| 58 | G C T A C G T A C G T A G C A T C A T G C T A G A G T C A C T G A C T G A G T C A C T G T C A G | ABR1(AP2EREBP)/colamp-ABR1-DAP-Seq(GSE60143)/Homer | 1e-4 | -1.066e+01 | 0.0004 | 302.0 | 27.13% | 9561.6 | 21.92% | motif file (matrix) | svg |
| 59 | A G C T C T G A A G T C A C T G A G C T T G C A C G T A A G T C | ATAF1(NAC)/col-ATAF1-DAP-Seq(GSE60143)/Homer | 1e-4 | -1.046e+01 | 0.0005 | 492.0 | 44.20% | 16686.5 | 38.25% | motif file (matrix) | svg |
| 60 | C G T A C T A G A C T G A C T G G A C T C T A G C A G T C T A G C A T G G A T C | KLF5(Zf)/LoVo-KLF5-ChIP-Seq(GSE49402)/Homer | 1e-4 | -1.043e+01 | 0.0005 | 415.0 | 37.29% | 13771.2 | 31.57% | motif file (matrix) | svg |
| 61 | C G T A T A G C T A G C T G C A A C T G C T A G C G T A C G T A T C A G G A C T | EHF(ETS)/LoVo-EHF-ChIP-Seq(GSE49402)/Homer | 1e-4 | -1.027e+01 | 0.0006 | 317.0 | 28.48% | 10158.2 | 23.29% | motif file (matrix) | svg |
| 62 | C T G A C T G A G C A T G A C T A G T C T C G A C T A G C G T A A C G T G A T C A G C T T C A G | GATA4(C2C2gata)/col-GATA4-DAP-Seq(GSE60143)/Homer | 1e-4 | -1.026e+01 | 0.0006 | 162.0 | 14.56% | 4653.9 | 10.67% | motif file (matrix) | svg |
| 63 | C A T G T G A C C G T A A G T C T A C G G C A T A C T G G T C A A T G C A G T C | bHLHE41(bHLH)/proB-Bhlhe41-ChIP-Seq(GSE93764)/Homer | 1e-4 | -1.017e+01 | 0.0006 | 317.0 | 28.48% | 10170.8 | 23.31% | motif file (matrix) | svg |
| 64 | C T A G C G A T C T A G C G A T C T G A G C A T T C A G C A G T A G T C C G T A A G T C A C T G A C G T A C T G G T C A | BIM1(bHLH)/colamp-BIM1-DAP-Seq(GSE60143)/Homer | 1e-4 | -9.948e+00 | 0.0007 | 43.0 | 3.86% | 870.2 | 1.99% | motif file (matrix) | svg |
| 65 | C G T A G C A T G A C T C T A G A C G T G T C A A G T C A C T G C T A G G C T A G T A C G C T A | SPL5(SBP)/colamp-SPL5-DAP-Seq(GSE60143)/Homer | 1e-4 | -9.881e+00 | 0.0008 | 103.0 | 9.25% | 2714.0 | 6.22% | motif file (matrix) | svg |
| 66 | C G T A G C A T C A G T C T A G A G T C A C T G A C T G G T A C A C T G A T C G | ERF115(AP2EREBP)/colamp-ERF115-DAP-Seq(GSE60143)/Homer | 1e-4 | -9.838e+00 | 0.0008 | 423.0 | 38.01% | 14158.9 | 32.46% | motif file (matrix) | svg |
| 67 | C A T G G T A C C G T A A G T C C T A G A C G T A C T G G T A C A G T C A G C T | bHLHE40(bHLH)/HepG2-BHLHE40-ChIP-Seq(GSE31477)/Homer | 1e-4 | -9.838e+00 | 0.0008 | 97.0 | 8.72% | 2523.4 | 5.78% | motif file (matrix) | svg |
| 68 | T C G A T A G C T G A C C T G A A G T C A C T G G A C T C A T G | c-Myc(bHLH)/LNCAP-cMyc-ChIP-Seq(Unpublished)/Homer | 1e-4 | -9.805e+00 | 0.0008 | 162.0 | 14.56% | 4695.7 | 10.76% | motif file (matrix) | svg |
| 69 | T C A G A C G T A G T C T C G A A G T C T C A G G C A T C T A G C T A G A G C T | Usf2(bHLH)/C2C12-Usf2-ChIP-Seq(GSE36030)/Homer | 1e-4 | -9.726e+00 | 0.0009 | 90.0 | 8.09% | 2306.9 | 5.29% | motif file (matrix) | svg |
| 70 | T G A C C T A G T C A G G T C A C G T A T C A G C G A T T C A G T C G A T G C A C T G A T A G C | PU.1-IRF(ETS:IRF)/Bcell-PU.1-ChIP-Seq(GSE21512)/Homer | 1e-4 | -9.656e+00 | 0.0009 | 295.0 | 26.50% | 9433.1 | 21.62% | motif file (matrix) | svg |
| 71 | G C A T A T C G C A T G G T A C G C T A A G T C T C A G T G A C G T C A T G C A | Arnt:Ahr(bHLH)/MCF7-Arnt-ChIP-Seq(Lo\_et\_al.)/Homer | 1e-4 | -9.581e+00 | 0.0010 | 190.0 | 17.07% | 5687.9 | 13.04% | motif file (matrix) | svg |
| 72 | C G A T C T G A C T A G G T A C A C T G A G T C C T A G A G T C | DPL-1(E2F)/cElegans-Adult-ChIP-Seq(modEncode)/Homer | 1e-4 | -9.568e+00 | 0.0010 | 294.0 | 26.42% | 9408.3 | 21.57% | motif file (matrix) | svg |
| 73 | G C T A C G T A C G T A G C A T C A T G C T A G A G T C A C T G T A C G A G T C C A T G T A C G | RAP26(AP2EREBP)/colamp-RAP26-DAP-Seq(GSE60143)/Homer | 1e-4 | -9.472e+00 | 0.0010 | 355.0 | 31.90% | 11666.4 | 26.74% | motif file (matrix) | svg |
| 74 | G C T A C T G A C T A G C G A T C A G T T C G A A G T C A C T G A C G T T C A G G C A T G C T A | NAP(NAC)/col-NAP-DAP-Seq(GSE60143)/Homer | 1e-4 | -9.463e+00 | 0.0010 | 167.0 | 15.00% | 4900.3 | 11.23% | motif file (matrix) | svg |
| 75 | G T A C A C T G A G T C A G T C C T A G G A T C G T A C C T G A | CRF4(AP2EREBP)/colamp-CRF4-DAP-Seq(GSE60143)/Homer | 1e-4 | -9.316e+00 | 0.0012 | 241.0 | 21.65% | 7523.9 | 17.25% | motif file (matrix) | svg |
| 76 | A T G C G T A C A C T G A G T C A G T C A C T G G A T C G T C A C G T A C G A T G C A T C G A T | RRTF1(AP2EREBP)/colamp-RRTF1-DAP-Seq(GSE60143)/Homer | 1e-4 | -9.282e+00 | 0.0012 | 78.0 | 7.01% | 1954.1 | 4.48% | motif file (matrix) | svg |
| 77 | C G T A C G A T C G A T G C A T C G A T T C A G A G T C A C T G C T A G A G T C A C G T C T G A | At5g08750(C3H)/col-At5g08750-DAP-Seq(GSE60143)/Homer | 1e-4 | -9.211e+00 | 0.0013 | 127.0 | 11.41% | 3560.5 | 8.16% | motif file (matrix) | svg |
| 78 | C T G A C G A T A C G T A C G T A C G T C T G A A G T C A C T G C G T A A C G T | ARF16(ARF)/col-ARF16-DAP-Seq(GSE60143)/Homer | 1e-3 | -9.168e+00 | 0.0013 | 32.0 | 2.88% | 597.1 | 1.37% | motif file (matrix) | svg |
| 79 | A G C T C G A T T A C G A T C G A T C G C A G T A G T C A G T C A C T G T A G C | HINFP(Zf)/K562-HINFP.eGFP-ChIP-Seq(Encode)/Homer | 1e-3 | -9.128e+00 | 0.0014 | 154.0 | 13.84% | 4485.8 | 10.28% | motif file (matrix) | svg |
| 80 | A T G C G T A C A C T G A G T C A G T C A C T G A G T C G T A C | ERF73(AP2EREBP)/col-ERF73-DAP-Seq(GSE60143)/Homer | 1e-3 | -9.105e+00 | 0.0014 | 284.0 | 25.52% | 9104.9 | 20.87% | motif file (matrix) | svg |
| 81 | C T A G G T A C A C G T A C G T A T C G G C A T A G C T A G C T A G C T G C A T G A C T C G T A G T C A A C T G G A C T | VND6(NAC)/col-VND6-DAP-Seq(GSE60143)/Homer | 1e-3 | -9.037e+00 | 0.0015 | 215.0 | 19.32% | 6628.7 | 15.20% | motif file (matrix) | svg |
| 82 | G T A C A C T G A T G C A G T C C T A G G A T C G T A C C T G A G A C T G C A T C G A T G A C T | RAP212(AP2EREBP)/col-RAP212-DAP-Seq(GSE60143)/Homer | 1e-3 | -9.021e+00 | 0.0015 | 256.0 | 23.00% | 8099.7 | 18.57% | motif file (matrix) | svg |
| 83 | C G T A C G T A C G A T A C T G C A G T A G T C A C T G A C T G A G C T A C T G | DREB19(AP2EREBP)/colamp-DREB19-DAP-Seq(GSE60143)/Homer | 1e-3 | -8.769e+00 | 0.0019 | 141.0 | 12.67% | 4074.1 | 9.34% | motif file (matrix) | svg |
| 84 | C T A G T A G C A T G C C T A G A G T C A G T C C T A G G A C T G A C T G C T A | CRF10(AP2EREBP)/col100-CRF10-DAP-Seq(GSE60143)/Homer | 1e-3 | -8.728e+00 | 0.0019 | 376.0 | 33.78% | 12558.2 | 28.79% | motif file (matrix) | svg |
| 85 | T A C G T C A G A G C T A T G C C G T A A G T C T C A G A C G T A C T G T C G A | USF1(bHLH)/GM12878-Usf1-ChIP-Seq(GSE32465)/Homer | 1e-3 | -8.627e+00 | 0.0021 | 128.0 | 11.50% | 3645.4 | 8.36% | motif file (matrix) | svg |
| 86 | C G T A C G A T C T A G T G C A A G C T A C T G C G T A A G T C C T A G C G A T T G A C C T G A A C G T G T A C G C T A | TGA5(bZIP)/col-TGA5-DAP-Seq(GSE60143)/Homer | 1e-3 | -8.517e+00 | 0.0023 | 33.0 | 2.96% | 647.8 | 1.49% | motif file (matrix) | svg |
| 87 | C A T G G A T C C T G A G T A C C T A G C T G A G C T A G C A T G A T C G A T C A G T C C T A G C G T A C A T G C T A G | AIL7(AP2EREBP)/colamp-AIL7-DAP-Seq(GSE60143)/Homer | 1e-3 | -8.514e+00 | 0.0023 | 120.0 | 10.78% | 3385.9 | 7.76% | motif file (matrix) | svg |
| 88 | C G T A C A G T G T A C A T G C C T A G C G T A A C G T A G T C T C G A T C A G | GATA19(C2C2gata)/colamp-GATA19-DAP-Seq(GSE60143)/Homer | 1e-3 | -8.424e+00 | 0.0025 | 24.0 | 2.16% | 411.4 | 0.94% | motif file (matrix) | svg |
| 89 | T C A G A G C T A T G C C G T A A G C T T C A G C A G T A C T G C T G A A G T C | MITF(bHLH)/MastCells-MITF-ChIP-Seq(GSE48085)/Homer | 1e-3 | -8.423e+00 | 0.0025 | 218.0 | 19.59% | 6808.4 | 15.61% | motif file (matrix) | svg |
| 90 | C T G A C A G T C T G A A G T C C T A G G A C T A T C G G T A C | HIF-1b(HLH)/T47D-HIF1b-ChIP-Seq(GSE59937)/Homer | 1e-3 | -8.334e+00 | 0.0027 | 285.0 | 25.61% | 9245.5 | 21.19% | motif file (matrix) | svg |
| 91 | A G C T A G C T T A G C A T C G A G T C A C T G A T G C A T C G T C G A C T G A T C G A C T G A | E2F(E2F)/Hela-CellCycle-Expression/Homer | 1e-3 | -8.275e+00 | 0.0028 | 33.0 | 2.96% | 656.3 | 1.50% | motif file (matrix) | svg |
| 92 | T C G A G A C T A T C G C G T A A G T C C T A G G C A T G T A C C T G A A C G T G A T C G C T A | TGA4(bZIP)/colamp-TGA4-DAP-Seq(GSE60143)/Homer | 1e-3 | -8.247e+00 | 0.0028 | 71.0 | 6.38% | 1799.2 | 4.12% | motif file (matrix) | svg |
| 93 | T A G C C G A T A C G T A G C T A G C T A G T C A T G C A G T C A C T G A T G C A T G C G C T A | E2F7(E2F)/Hela-E2F7-ChIP-Seq(GSE32673)/Homer | 1e-3 | -8.225e+00 | 0.0029 | 63.0 | 5.66% | 1549.4 | 3.55% | motif file (matrix) | svg |
| 94 | C G A T C G T A C G A T C G A T A G T C C T G A G A T C C T G A G A T C C T A G G C A T C T A G G A C T T C A G G C T A | At4g36780(BZR)/col-At4g36780-DAP-Seq(GSE60143)/Homer | 1e-3 | -8.213e+00 | 0.0029 | 86.0 | 7.73% | 2283.2 | 5.23% | motif file (matrix) | svg |
| 95 | C T A G G T A C A G T C T G C A A G T C C T G A A G T C A G T C A G T C G C T A | Klf4(Zf)/mES-Klf4-ChIP-Seq(GSE11431)/Homer | 1e-3 | -8.212e+00 | 0.0029 | 139.0 | 12.49% | 4060.0 | 9.31% | motif file (matrix) | svg |
| 96 | C G A T G A T C T A C G C T G A G C T A C G T A G C A T A G T C C T A G C G T A G C A T C G A T | AT2G15740(C2H2)/col-AT2G15740-DAP-Seq(GSE60143)/Homer | 1e-3 | -8.181e+00 | 0.0029 | 139.0 | 12.49% | 4062.6 | 9.31% | motif file (matrix) | svg |
| 97 | A G T C T C G A T C A G C G T A G C A T G A T C A G C T T C A G C T G A C G T A | GATA12(C2C2gata)/col-GATA12-DAP-Seq(GSE60143)/Homer | 1e-3 | -8.152e+00 | 0.0030 | 131.0 | 11.77% | 3791.5 | 8.69% | motif file (matrix) | svg |
| 98 | G A T C G T A C C T G A A G T C A G T C A C T G G C T A G T A C G T C A G C A T G C A T C G A T | DEAR2(AP2EREBP)/colamp-DEAR2-DAP-Seq(GSE60143)/Homer | 1e-3 | -8.078e+00 | 0.0031 | 248.0 | 22.28% | 7932.1 | 18.18% | motif file (matrix) | svg |
| 99 | C A T G C T G A A G T C A C T G A C T G A G C T A C T G A T C G | ESE3(AP2EREBP)/col-ESE3-DAP-Seq(GSE60143)/Homer | 1e-3 | -8.037e+00 | 0.0032 | 377.0 | 33.87% | 12703.7 | 29.12% | motif file (matrix) | svg |
| 100 | T C A G C T G A C G T A C G T A T A C G G C A T C T A G C T G A C G T A C G T A T A C G G A C T | IRF1(IRF)/PBMC-IRF1-ChIP-Seq(GSE43036)/Homer | 1e-3 | -7.992e+00 | 0.0034 | 37.0 | 3.32% | 781.0 | 1.79% | motif file (matrix) | svg |
| 101 | A G T C A C T G A C G T C A T G C T G A G C T A C G A T C G A T G A C T G A C T G T C A G T A C A C T G A C T G G A T C | ANAC042(NAC)/col-ANAC042-DAP-Seq(GSE60143)/Homer | 1e-3 | -7.953e+00 | 0.0035 | 200.0 | 17.97% | 6223.0 | 14.27% | motif file (matrix) | svg |
| 102 | G A C T A G T C C T G A A G T C A G T C A C T G C T G A A G T C G C T A G C A T G T A C C G A T G C A T G A C T C G A T | CBF2(AP2EREBP)/colamp-CBF2-DAP-Seq(GSE60143)/Homer | 1e-3 | -7.908e+00 | 0.0036 | 116.0 | 10.42% | 3304.6 | 7.58% | motif file (matrix) | svg |
| 103 | A T G C A T G C T A G C C G T A T G A C A C T G G C A T C T A G G T A C A G C T | Pho2(bHLH)/Yeast-Pho2-ChIP-Seq(GSE29506)/Homer | 1e-3 | -7.905e+00 | 0.0036 | 127.0 | 11.41% | 3678.7 | 8.43% | motif file (matrix) | svg |
| 104 | G C A T C T A G A C T G A C G T C G T A A C T G A C T G C G A T C T A G T C G A T C G A G C T A | MYB40(MYB)/col-MYB40-DAP-Seq(GSE60143)/Homer | 1e-3 | -7.837e+00 | 0.0038 | 78.0 | 7.01% | 2051.1 | 4.70% | motif file (matrix) | svg |
| 105 | T A G C C T A G T C G A G A C T A C T G C T G A A G T C T C A G G C A T T G A C C T G A A G C T | Atf7(bZIP)/3T3L1-Atf7-ChIP-Seq(GSE56872)/Homer | 1e-3 | -7.672e+00 | 0.0044 | 109.0 | 9.79% | 3089.5 | 7.08% | motif file (matrix) | svg |
| 106 | G C T A G A C T A G T C T C G A T C A G T C G A A C G T A G T C G A C T T C A G | GATA14(C2C2gata)/col-GATA14-DAP-Seq(GSE60143)/Homer | 1e-3 | -7.608e+00 | 0.0047 | 146.0 | 13.12% | 4362.8 | 10.00% | motif file (matrix) | svg |
| 107 | A T G C T C A G T C G A G C A T A C T G C G T A A G T C T C A G G A C T T G A C C G T A A G C T | Atf2(bZIP)/3T3L1-Atf2-ChIP-Seq(GSE56872)/Homer | 1e-3 | -7.597e+00 | 0.0047 | 84.0 | 7.55% | 2264.7 | 5.19% | motif file (matrix) | svg |
| 108 | C T A G G C A T G A T C C G T A A G T C T C A G G A C T C T A G | CLOCK(bHLH)/Liver-Clock-ChIP-Seq(GSE39860)/Homer | 1e-3 | -7.589e+00 | 0.0047 | 143.0 | 12.85% | 4260.1 | 9.77% | motif file (matrix) | svg |
| 109 | A C T G G T A C A G T C C T G A A T G C A C T G A G C T T A C G | E-box/Arabidopsis-Promoters/Homer | 1e-3 | -7.574e+00 | 0.0047 | 127.0 | 11.41% | 3710.1 | 8.50% | motif file (matrix) | svg |
| 110 | G A C T A C T G C A G T A G T C A C T G A C T G A G C T A C T G C T A G G T C A | At1g77640(AP2EREBP)/col-At1g77640-DAP-Seq(GSE60143)/Homer | 1e-3 | -7.564e+00 | 0.0047 | 61.0 | 5.48% | 1527.4 | 3.50% | motif file (matrix) | svg |
| 111 | A T G C G A T C G A C T A G C T C G A T C G A T G T C A C G A T T C G A A T C G T A G C T A G C | TATA-Box(TBP)/Promoter/Homer | 1e-3 | -7.481e+00 | 0.0050 | 227.0 | 20.40% | 7252.1 | 16.62% | motif file (matrix) | svg |
| 112 | C A T G A G T C G T A C A C T G A T G C A G T C C A T G G A T C G A T C C T G A | ERF5(AP2EREBP)/colamp-ERF5-DAP-Seq(GSE60143)/Homer | 1e-3 | -7.467e+00 | 0.0051 | 197.0 | 17.70% | 6176.5 | 14.16% | motif file (matrix) | svg |
| 113 | G T C A T A C G C G A T C T A G C T G A G C A T C G A T C A T G T C G A A G T C C G T A A G T C A C T G A C G T A C T G | bHLH34(bHLH)/colamp-bHLH34-DAP-Seq(GSE60143)/Homer | 1e-3 | -7.373e+00 | 0.0055 | 78.0 | 7.01% | 2085.1 | 4.78% | motif file (matrix) | svg |
| 114 | C G A T A G C T T G C A A C T G A G T C T G A C C T A G G T A C A G T C C G T A G C A T G C A T | ERF13(AP2EREBP)/colamp-ERF13-DAP-Seq(GSE60143)/Homer | 1e-3 | -7.359e+00 | 0.0056 | 332.0 | 29.83% | 11128.6 | 25.51% | motif file (matrix) | svg |
| 115 | C A G T T A G C A G T C C A T G C A G T C A T G C G A T C G A T G A C T C G A T A T C G G T A C A C T G A T C G G T A C | LBD13(LOBAS2)/colamp-LBD13-DAP-Seq(GSE60143)/Homer | 1e-3 | -7.242e+00 | 0.0062 | 255.0 | 22.91% | 8303.4 | 19.03% | motif file (matrix) | svg |
| 116 | G C A T T A G C G T A C C A T G C T G A G C A T G C A T G C A T G A C T G C A T G A C T G T A C A C T G A T C G C G T A | LBD2(LOBAS2)/colamp-LBD2-DAP-Seq(GSE60143)/Homer | 1e-3 | -7.220e+00 | 0.0063 | 80.0 | 7.19% | 2162.6 | 4.96% | motif file (matrix) | svg |
| 117 | G T A C A C T G A T G C T G A C C T A G G A C T G T A C C G T A G C A T G C A T | ERF8(AP2EREBP)/colamp-ERF8-DAP-Seq(GSE60143)/Homer | 1e-3 | -7.078e+00 | 0.0072 | 313.0 | 28.12% | 10467.0 | 23.99% | motif file (matrix) | svg |
| 118 | C T A G T C A G C A G T T C A G A C T G A C T G G A T C C T A G A C T G C T A G T C A G A T G C | KLF14(Zf)/HEK293-KLF14.GFP-ChIP-Seq(GSE58341)/Homer | 1e-3 | -7.024e+00 | 0.0075 | 502.0 | 45.10% | 17641.1 | 40.44% | motif file (matrix) | svg |
| 119 | A G T C C A G T T C A G A T G C A G T C C G A T C G T A G T C A G A T C G C A T | BOS1(MYB)/col-BOS1-DAP-Seq(GSE60143)/Homer | 1e-3 | -7.015e+00 | 0.0075 | 184.0 | 16.53% | 5769.9 | 13.23% | motif file (matrix) | svg |
| 120 | T A C G T C G A G A C T A C T G C T G A A G T C T C A G G A C T T G A C C T G A | Atf1(bZIP)/K562-ATF1-ChIP-Seq(GSE31477)/Homer | 1e-3 | -6.995e+00 | 0.0076 | 142.0 | 12.76% | 4289.1 | 9.83% | motif file (matrix) | svg |
| 121 | C A G T A T C G C T G A A G T C T C A G C A G T T A G C C T G A A T G C T A C G | FEA4(bZIP)/Corn-FEA4-ChIP-Seq(GSE61954)/Homer | 1e-3 | -6.961e+00 | 0.0078 | 237.0 | 21.29% | 7687.0 | 17.62% | motif file (matrix) | svg |
| 122 | A G T C A C G T A C G T T C A G G C T A C G T A G C T A G C A T C G A T A G T C C G T A G T C A A C T G G A C T G C T A | SMB(NAC)/colamp-SMB-DAP-Seq(GSE60143)/Homer | 1e-3 | -6.936e+00 | 0.0079 | 170.0 | 15.27% | 5281.3 | 12.11% | motif file (matrix) | svg |
| 123 | T A G C G T A C C T A G A T C G C T G A C G T A G C T A G C A T A C G T T G A C G T A C A C T G T A C G G T C A C T A G | ASL18(LOBAS2)/colamp-ASL18-DAP-Seq(GSE60143)/Homer | 1e-2 | -6.860e+00 | 0.0085 | 390.0 | 35.04% | 13391.4 | 30.70% | motif file (matrix) | svg |
| 124 | T C G A T C A G C G A T C A G T T C G A A G T C A C T G A C G T T C A G G C A T | NAM(NAC)/col-NAM-DAP-Seq(GSE60143)/Homer | 1e-2 | -6.812e+00 | 0.0088 | 217.0 | 19.50% | 6982.3 | 16.01% | motif file (matrix) | svg |
| 125 | G C A T A C T G C T A G A G T C A C T G A C T G A G T C A C G T | ERF105(AP2EREBP)/colamp-ERF105-DAP-Seq(GSE60143)/Homer | 1e-2 | -6.803e+00 | 0.0088 | 398.0 | 35.76% | 13704.7 | 31.42% | motif file (matrix) | svg |
| 126 | T C G A G C A T A C T G C T G A A G T C T C A G G A C T G T A C C G T A A G C T A G T C G A T C | c-Jun-CRE(bZIP)/K562-cJun-ChIP-Seq(GSE31477)/Homer | 1e-2 | -6.775e+00 | 0.0090 | 73.0 | 6.56% | 1967.1 | 4.51% | motif file (matrix) | svg |
| 127 | T C G A T G A C G C A T A G C T C A G T G A T C G C T A G A T C G A C T A C G T G C A T A G T C | PRDM1(Zf)/Hela-PRDM1-ChIP-Seq(GSE31477)/Homer | 1e-2 | -6.700e+00 | 0.0096 | 114.0 | 10.24% | 3351.7 | 7.68% | motif file (matrix) | svg |
| 128 | C G A T C G T A A G T C A C G T A C G T T C A G C G T A C G T A G C A T G C A T G C A T A G T C C G T A G T C A A C T G | VND2(NAC)/col-VND2-DAP-Seq(GSE60143)/Homer | 1e-2 | -6.630e+00 | 0.0102 | 172.0 | 15.45% | 5390.3 | 12.36% | motif file (matrix) | svg |
| 129 | T C G A C T G A T A G C C T A G A C T G G T C A A C G T A G C T C G T A C T A G T A C G A C G T | bcd(Homeobox)/Embryo-Bcd-ChIP-Seq(GSE86966)/Homer | 1e-2 | -6.565e+00 | 0.0109 | 195.0 | 17.52% | 6222.5 | 14.26% | motif file (matrix) | svg |
| 130 | C A T G G C A T C G A T C G T A C G T A G A T C A G T C A G C T C G T A C G T A A C G T A G T C C G T A C G T A G C A T | DUX4(Homeobox)/Myoblasts-DUX4.V5-ChIP-Seq(GSE75791)/Homer | 1e-2 | -6.442e+00 | 0.0122 | 10.0 | 0.90% | 124.6 | 0.29% | motif file (matrix) | svg |
| 131 | G T A C C A T G T A G C A G T C C T A G C A T G C T G A C G T A G C A T G C A T A C G T G C A T G T A C A C T G A T C G | LOB(LOBAS2)/col-LOB-DAP-Seq(GSE60143)/Homer | 1e-2 | -6.410e+00 | 0.0125 | 96.0 | 8.63% | 2765.8 | 6.34% | motif file (matrix) | svg |
| 132 | G C T A G C A T G C A T C A G T A C G T G C A T G T C A G T A C G A T C T A C G | AT5G47660(Trihelix)/colamp-AT5G47660-DAP-Seq(GSE60143)/Homer | 1e-2 | -6.382e+00 | 0.0127 | 204.0 | 18.33% | 6572.6 | 15.07% | motif file (matrix) | svg |
| 133 | C T G A T G C A A G T C A C T G A C G T T C A G C G A T G C A T G C A T G A T C G C A T G A T C G T C A A G T C A C T G | ANAC094(NAC)/col-ANAC094-DAP-Seq(GSE60143)/Homer | 1e-2 | -6.304e+00 | 0.0137 | 108.0 | 9.70% | 3185.3 | 7.30% | motif file (matrix) | svg |
| 134 | C G T A G C A T C A T G C T A G A G T C A C T G A T C G G T A C A C T G T C A G | At2g33710(AP2EREBP)/colamp-At2g33710-DAP-Seq(GSE60143)/Homer | 1e-2 | -6.296e+00 | 0.0137 | 450.0 | 40.43% | 15781.2 | 36.18% | motif file (matrix) | svg |
| 135 | T A G C T G A C C A T G C T G A C G T A C G T A G C T A G C A T G C A T G A C T G T A C A C T G A T C G C G T A C T A G | CDM1(C3H)/colamp-CDM1-DAP-Seq(GSE60143)/Homer | 1e-2 | -6.295e+00 | 0.0137 | 30.0 | 2.70% | 652.5 | 1.50% | motif file (matrix) | svg |
| 136 | G T A C A C T G A C G T T C A G G C A T C G T A C G A T G C A T C G T A A G T C C G T A T G A C C A T G G A C T G C T A | ANAC083(NAC)/col-ANAC083-DAP-Seq(GSE60143)/Homer | 1e-2 | -6.193e+00 | 0.0149 | 128.0 | 11.50% | 3890.1 | 8.92% | motif file (matrix) | svg |
| 137 | A G C T C T A G G A T C A G T C C T A G C T G A A G T C G C T A G C A T T G C A | CBF1(AP2EREBP)/colamp-CBF1-DAP-Seq(GSE60143)/Homer | 1e-2 | -6.153e+00 | 0.0154 | 159.0 | 14.29% | 4989.0 | 11.44% | motif file (matrix) | svg |
| 138 | C T A G A G T C A G T C A C T G C G T A A G T C C T G A G A C T | DDF1(AP2EREBP)/col-DDF1-DAP-Seq(GSE60143)/Homer | 1e-2 | -6.116e+00 | 0.0159 | 93.0 | 8.36% | 2692.0 | 6.17% | motif file (matrix) | svg |
| 139 | G C T A C G A T C T A G T C G A A G C T A C T G C G T A A G T C C T A G A C G T G T A C C T G A A G C T G A T C C G T A | TGA3(bZIP)/colamp-TGA3-DAP-Seq(GSE60143)/Homer | 1e-2 | -6.044e+00 | 0.0170 | 22.0 | 1.98% | 435.4 | 1.00% | motif file (matrix) | svg |
| 140 | C G T A C G T A G C A T G A C T A G T C C T G A C T A G C G T A A C G T A G T C A G C T T C A G | GATA11(C2C2gata)/col-GATA11-DAP-Seq(GSE60143)/Homer | 1e-2 | -6.032e+00 | 0.0170 | 92.0 | 8.27% | 2665.7 | 6.11% | motif file (matrix) | svg |
| 141 | C T A G C T A G C G T A C G T A T A C G C G A T C T A G C T G A C T G A C G T A T A C G G A C T | PU.1:IRF8(ETS:IRF)/pDC-Irf8-ChIP-Seq(GSE66899)/Homer | 1e-2 | -6.021e+00 | 0.0171 | 49.0 | 4.40% | 1247.6 | 2.86% | motif file (matrix) | svg |
| 142 | T C G A A C G T A C G T C T G A G A T C T C A G G A C T G T C A C G T A A G C T G T C A C T A G A G C T A C G T T C G A | NFIL3(bZIP)/HepG2-NFIL3-ChIP-Seq(Encode)/Homer | 1e-2 | -6.020e+00 | 0.0171 | 99.0 | 8.89% | 2905.5 | 6.66% | motif file (matrix) | svg |
| 143 | C G T A C A G T C A T G G T A C A G T C C G T A A T G C A T C G A C G T T A C G C G T A G A T C G T A C C G T A C G T A | AREB3(bZIP)/col-AREB3-DAP-Seq(GSE60143)/Homer | 1e-2 | -6.006e+00 | 0.0171 | 75.0 | 6.74% | 2096.0 | 4.80% | motif file (matrix) | svg |
| 144 | C T G A G C T A G C A T A G C T A C T G A C G T C G T A A G T C A C T G C T A G G C T A G A T C | bHLH28(bHLH)/col-bHLH28-DAP-Seq(GSE60143)/Homer | 1e-2 | -5.961e+00 | 0.0178 | 59.0 | 5.30% | 1572.9 | 3.61% | motif file (matrix) | svg |
| 145 | T A G C A T G C C G T A A G T C C T A G G A C T T A C G T C A G A G C T G C T A | PIF7(bHLH)/col-PIF7-DAP-Seq(GSE60143)/Homer | 1e-2 | -5.893e+00 | 0.0189 | 59.0 | 5.30% | 1577.6 | 3.62% | motif file (matrix) | svg |
| 146 | A C G T C T G A G A T C A T C G G A C T T C A G G T A C T A G C | HIF-1a(bHLH)/MCF7-HIF1a-ChIP-Seq(GSE28352)/Homer | 1e-2 | -5.877e+00 | 0.0191 | 77.0 | 6.92% | 2173.8 | 4.98% | motif file (matrix) | svg |
| 147 | A C T G A C T G A G T C A C T G A C T G A G T C A C G T C T A G | ERF1(AP2EREBP)/colamp-ERF1-DAP-Seq(GSE60143)/Homer | 1e-2 | -5.852e+00 | 0.0194 | 247.0 | 22.19% | 8223.7 | 18.85% | motif file (matrix) | svg |
| 148 | C G A T C T G A A G T C A C G T A C G T T C A G C G T A C G T A C G T A G C A T C G A T A G T C C G T A G T C A A C T G | VND4(NAC)/colamp-VND4-DAP-Seq(GSE60143)/Homer | 1e-2 | -5.841e+00 | 0.0195 | 131.0 | 11.77% | 4037.0 | 9.25% | motif file (matrix) | svg |
| 149 | T C A G G A C T A C G T C G T A A C T G A C T G A C T G A G T C C G T A G T C A | TBP3(MYBrelated)/col-TBP3-DAP-Seq(GSE60143)/Homer | 1e-2 | -5.805e+00 | 0.0201 | 173.0 | 15.54% | 5535.6 | 12.69% | motif file (matrix) | svg |
| 150 | C G T A G A C T G A C T C T G A C T G A C G T A G T A C A T G C T A C G C T A G | Unknown3/Arabidopsis-Promoters/Homer | 1e-2 | -5.763e+00 | 0.0208 | 49.0 | 4.40% | 1264.2 | 2.90% | motif file (matrix) | svg |
| 151 | C G T A G A C T C A T G C T A G A G T C A C T G A C T G G T A C C A T G T A C G | ERF3(AP2EREBP)/colamp-ERF3-DAP-Seq(GSE60143)/Homer | 1e-2 | -5.701e+00 | 0.0220 | 235.0 | 21.11% | 7806.9 | 17.90% | motif file (matrix) | svg |
| 152 | C T G A G A C T C A T G C T A G A G T C A C T G A C T G A G T C A C T G T C A G | ERF11(AP2EREBP)/col-ERF11-DAP-Seq(GSE60143)/Homer | 1e-2 | -5.698e+00 | 0.0220 | 281.0 | 25.25% | 9508.1 | 21.80% | motif file (matrix) | svg |
| 153 | G A T C C T G A G A T C G A T C C T A G G C T A A G T C C T G A G C T A C G T A | At4g16750(AP2EREBP)/col-At4g16750-DAP-Seq(GSE60143)/Homer | 1e-2 | -5.665e+00 | 0.0225 | 166.0 | 14.91% | 5303.9 | 12.16% | motif file (matrix) | svg |
| 154 | A G T C G A C T C A G T G T A C A G T C A T C G T C A G A C T G G T C A C G T A | Stat3(Stat)/mES-Stat3-ChIP-Seq(GSE11431)/Homer | 1e-2 | -5.651e+00 | 0.0227 | 131.0 | 11.77% | 4059.3 | 9.31% | motif file (matrix) | svg |
| 155 | G C A T T G A C C T G A A G T C A G T C A C T G G T C A A G T C G C T A G A C T G C T A C T G A | DREB2(AP2EREBP)/col-DREB2-DAP-Seq(GSE60143)/Homer | 1e-2 | -5.637e+00 | 0.0229 | 110.0 | 9.88% | 3325.1 | 7.62% | motif file (matrix) | svg |
| 156 | C G A T T C G A A G T C A C G T A C G T T A C G C G T A G C T A G C T A C G A T G C A T A T G C C G T A G T C A A C T G | ANAC071(NAC)/col-ANAC071-DAP-Seq(GSE60143)/Homer | 1e-2 | -5.630e+00 | 0.0229 | 154.0 | 13.84% | 4878.8 | 11.18% | motif file (matrix) | svg |
| 157 | C G A T T C A G G T A C A C G T A C G T T C A G C G A T C G T A G T C A G C T A C G T A A G T C C G T A G T C A C A T G | ANAC057(NAC)/colamp-ANAC057-DAP-Seq(GSE60143)/Homer | 1e-2 | -5.586e+00 | 0.0238 | 139.0 | 12.49% | 4350.3 | 9.97% | motif file (matrix) | svg |
| 158 | G C T A C G T A C G T A G C A T C A T G C T A G G A T C A C T G T C A G G A T C A C T G T C A G | ERF4(AP2EREBP)/colamp-ERF4-DAP-Seq(GSE60143)/Homer | 1e-2 | -5.564e+00 | 0.0241 | 308.0 | 27.67% | 10540.0 | 24.16% | motif file (matrix) | svg |
| 159 | C T G A G C A T A C T G C T A G A G T C A C T G A C T G A G T C A C T G T C A G | AT4G18450(AP2EREBP)/col-AT4G18450-DAP-Seq(GSE60143)/Homer | 1e-2 | -5.553e+00 | 0.0242 | 179.0 | 16.08% | 5787.8 | 13.27% | motif file (matrix) | svg |
| 160 | G C T A C G T A C G T A G A C T C A T G C T A G G A T C A C T G T C A G G A T C A C T G T A C G | ERF9(AP2EREBP)/colamp-ERF9-DAP-Seq(GSE60143)/Homer | 1e-2 | -5.546e+00 | 0.0242 | 123.0 | 11.05% | 3790.4 | 8.69% | motif file (matrix) | svg |
| 161 | C G T A T G A C T A G C T G C A A C T G A C T G C G T A C G T A T C A G G A C T | ELF3(ETS)/PDAC-ELF3-ChIP-Seq(GSE64557)/Homer | 1e-2 | -5.521e+00 | 0.0247 | 171.0 | 15.36% | 5503.3 | 12.62% | motif file (matrix) | svg |
| 162 | G C T A A G T C T A C G T G C A A T C G T C A G G C T A T C G A T C A G A G C T | ELF5(ETS)/T47D-ELF5-ChIP-Seq(GSE30407)/Homer | 1e-2 | -5.515e+00 | 0.0247 | 182.0 | 16.35% | 5901.1 | 13.53% | motif file (matrix) | svg |
| 163 | A C T G C G T A C G T A A C G T G T A C G A C T C G T A C G A T C T G A C T G A | AT1G49560(G2like)/colamp-AT1G49560-DAP-Seq(GSE60143)/Homer | 1e-2 | -5.503e+00 | 0.0248 | 250.0 | 22.46% | 8392.3 | 19.24% | motif file (matrix) | svg |
| 164 | G A C T G A T C C T G A A G T C A G T C A C T G C G T A A G T C G T A C G C T A G C A T C G A T | At1g19210(AP2EREBP)/colamp-At1g19210-DAP-Seq(GSE60143)/Homer | 1e-2 | -5.503e+00 | 0.0248 | 298.0 | 26.77% | 10177.6 | 23.33% | motif file (matrix) | svg |
| 165 | A C T G C T A G A G T C A C T G A C T G A G T C A C T G T A C G | ERF104(AP2EREBP)/col-ERF104-DAP-Seq(GSE60143)/Homer | 1e-2 | -5.421e+00 | 0.0266 | 334.0 | 30.01% | 11544.4 | 26.46% | motif file (matrix) | svg |
| 166 | G A T C A T G C A G T C C G T A A G T C A G T C A C T G G C T A A G T C C G T A | AT1G44830(AP2EREBP)/col-AT1G44830-DAP-Seq(GSE60143)/Homer | 1e-2 | -5.417e+00 | 0.0266 | 94.0 | 8.45% | 2795.3 | 6.41% | motif file (matrix) | svg |
| 167 | C A T G A C T G A G C T A T G C C G T A A T G C G T A C G A C T T A C G C T G A A C T G A C T G G C A T A T G C C T G A | THRb(NR)/HepG2-THRb.Flag-ChIP-Seq(Encode)/Homer | 1e-2 | -5.415e+00 | 0.0266 | 163.0 | 14.65% | 5230.1 | 11.99% | motif file (matrix) | svg |
| 168 | A G C T C T A G A G T C A G T C A C T G C G T A A G T C C T G A G C A T G C T A C T G A G C A T G C A T C G A T G C A T | CBF4(AP2EREBP)/colamp-CBF4-DAP-Seq(GSE60143)/Homer | 1e-2 | -5.371e+00 | 0.0275 | 155.0 | 13.93% | 4949.9 | 11.35% | motif file (matrix) | svg |
| 169 | C G T A C G A T C A G T C A T G C G A T G T A C C A T G A C T G G A C T C A T G | CEJ1(AP2EREBP)/col-CEJ1-DAP-Seq(GSE60143)/Homer | 1e-2 | -5.360e+00 | 0.0276 | 219.0 | 19.68% | 7274.0 | 16.67% | motif file (matrix) | svg |
| 170 | C G T A G C A T C G T A C G T A G C A T A C T G C G A T A G T C A C T G A C T G G A C T C T A G | AT1G71450(AP2EREBP)/col-AT1G71450-DAP-Seq(GSE60143)/Homer | 1e-2 | -5.347e+00 | 0.0278 | 368.0 | 33.06% | 12844.7 | 29.44% | motif file (matrix) | svg |
| 171 | C G T A G A C T C A T G C T A G A G T C A C T G C T A G A G T C C A T G C T A G | ERF7(AP2EREBP)/col-ERF7-DAP-Seq(GSE60143)/Homer | 1e-2 | -5.312e+00 | 0.0287 | 352.0 | 31.63% | 12245.9 | 28.07% | motif file (matrix) | svg |
| 172 | A C T G C A T G G C T A T C G A G C T A A G C T A G C T G T A C A G T C T G A C | NFkB-p65-Rel(RHD)/ThioMac-LPS-Expression(GSE23622)/Homer | 1e-2 | -5.237e+00 | 0.0307 | 18.0 | 1.62% | 354.7 | 0.81% | motif file (matrix) | svg |
| 173 | C T A G G C A T A C T G C T A G A G T C A C T G A C T G A G T C A C T G T C A G | ERF10(AP2EREBP)/col-ERF10-DAP-Seq(GSE60143)/Homer | 1e-2 | -5.227e+00 | 0.0309 | 240.0 | 21.56% | 8070.3 | 18.50% | motif file (matrix) | svg |
| 174 | C T G A A T C G A G C T A G C T A C G T T A G C C T G A T A C G C G A T A C G T G A C T A G T C | ISRE(IRF)/ThioMac-LPS-Expression(GSE23622)/Homer | 1e-2 | -5.197e+00 | 0.0316 | 17.0 | 1.53% | 328.1 | 0.75% | motif file (matrix) | svg |
| 175 | G C T A C G T A G A C T A G C T A C T G A C G T C G T A A G T C A C T G C T A G G C T A G A T C | SPL13(SBP)/col-SPL13-DAP-Seq(GSE60143)/Homer | 1e-2 | -5.160e+00 | 0.0326 | 26.0 | 2.34% | 587.0 | 1.35% | motif file (matrix) | svg |
| 176 | A C T G T C A G T C A G C T G A G C T A G C T A G C T A G T A C G T A C T A G C T G A C T C A G | Dorsal(RHD)/Embryo-dl-ChIP-Seq(GSE65441)/Homer | 1e-2 | -5.131e+00 | 0.0334 | 48.0 | 4.31% | 1276.9 | 2.93% | motif file (matrix) | svg |
| 177 | C G T A C G T A C G T A C G T A C G T A A C T G A G C T C T A G G T A C G C T A | AT1G69570(C2C2dof)/col-AT1G69570-DAP-Seq(GSE60143)/Homer | 1e-2 | -5.130e+00 | 0.0334 | 206.0 | 18.51% | 6835.1 | 15.67% | motif file (matrix) | svg |
| 178 | A G T C G A T C C T G A A G T C A G T C C A T G G T C A G A T C C G T A G A T C | DREB26(AP2EREBP)/col-DREB26-DAP-Seq(GSE60143)/Homer | 1e-2 | -5.128e+00 | 0.0334 | 75.0 | 6.74% | 2174.7 | 4.99% | motif file (matrix) | svg |
| 179 | T C G A A C G T A C T G C G T A A G T C C T A G A G C T T G A C | TGA10(bZIP)/colamp-TGA10-DAP-Seq(GSE60143)/Homer | 1e-2 | -5.118e+00 | 0.0334 | 118.0 | 10.60% | 3666.2 | 8.40% | motif file (matrix) | svg |
| 180 | G C T A T A G C A G C T A T C G G T C A C G T A G C T A A T G C G A T C C T G A | IRF4(IRF)/GM12878-IRF4-ChIP-Seq(GSE32465)/Homer | 1e-2 | -5.110e+00 | 0.0334 | 84.0 | 7.55% | 2483.2 | 5.69% | motif file (matrix) | svg |
| 181 | C A T G T A C G T A G C G A T C G A T C A T G C G T A C G A C T T C A G A T G C C G A T A T C G C A G T A C T G G T A C | Zic3(Zf)/mES-Zic3-ChIP-Seq(GSE37889)/Homer | 1e-2 | -5.069e+00 | 0.0346 | 188.0 | 16.89% | 6187.0 | 14.18% | motif file (matrix) | svg |
| 182 | G A C T C G A T C A T G G C T A G C A T C G T A A G T C G C T A G C A T C G A T T A C G G C A T C G T A C A G T G A T C | DMRT6(DM)/Testis-DMRT6-ChIP-Seq(GSE60440)/Homer | 1e-2 | -5.028e+00 | 0.0358 | 36.0 | 3.23% | 900.2 | 2.06% | motif file (matrix) | svg |
| 183 | G A C T G A C T G A T C C G T A G T A C A G T C G C A T C G T A G T A C G A T C G C A T G C T A | MYB74(MYB)/colamp-MYB74-DAP-Seq(GSE60143)/Homer | 1e-2 | -5.016e+00 | 0.0360 | 111.0 | 9.97% | 3432.8 | 7.87% | motif file (matrix) | svg |
| 184 | C A T G T G C A G A C T C A T G C G T A A G T C T C A G G C A T T G A C C G T A | bZIP50(bZIP)/colamp-bZIP50-DAP-Seq(GSE60143)/Homer | 1e-2 | -5.005e+00 | 0.0362 | 190.0 | 17.07% | 6270.8 | 14.37% | motif file (matrix) | svg |
| 185 | C A T G A G T C C T G A T G A C A T C G G A C T G T C A A G T C T A G C G A T C | HIF2a(bHLH)/785\_O-HIF2a-ChIP-Seq(GSE34871)/Homer | 1e-2 | -4.943e+00 | 0.0383 | 104.0 | 9.34% | 3195.2 | 7.32% | motif file (matrix) | svg |
| 186 | C G A T G C A T A G T C T C A G G A T C C G T A G A T C C T A G G C A T C T A G A G C T T C G A C G T A G C T A G C A T | At1g78700(BZR)/col-At1g78700-DAP-Seq(GSE60143)/Homer | 1e-2 | -4.877e+00 | 0.0407 | 78.0 | 7.01% | 2301.9 | 5.28% | motif file (matrix) | svg |
| 187 | G C T A T G A C G A T C C G A T G A C T A T G C C T G A A T C G G C A T A C G T | JGL(C2H2)/col-JGL-DAP-Seq(GSE60143)/Homer | 1e-2 | -4.856e+00 | 0.0414 | 280.0 | 25.16% | 9626.5 | 22.07% | motif file (matrix) | svg |
| 188 | C G T A C G T A C T A G C A G T G A C T C G T A C A T G C A T G C G A T C T G A C T G A C T G A | MS188(MYB)/colamp-MS188-DAP-Seq(GSE60143)/Homer | 1e-2 | -4.854e+00 | 0.0414 | 120.0 | 10.78% | 3770.7 | 8.64% | motif file (matrix) | svg |
| 189 | G A T C C T G A A G T C G T A C A C T G G C T A G A T C C T G A G C T A G C T A | At4g31060(AP2EREBP)/colamp-At4g31060-DAP-Seq(GSE60143)/Homer | 1e-2 | -4.852e+00 | 0.0414 | 91.0 | 8.18% | 2752.4 | 6.31% | motif file (matrix) | svg |
| 190 | C G T A C G T A C G T A C G T A A G T C A G T C C T A G A T C G C G A T G C T A | AT1G76870(Trihelix)/col-AT1G76870-DAP-Seq(GSE60143)/Homer | 1e-2 | -4.848e+00 | 0.0414 | 70.0 | 6.29% | 2031.2 | 4.66% | motif file (matrix) | svg |
| 191 | C G T A A C G T A C G T A C G T A C G T A G T C A G T C C T G A A G C T A G C T | NFAT(RHD)/Jurkat-NFATC1-ChIP-Seq(Jolma\_et\_al.)/Homer | 1e-2 | -4.836e+00 | 0.0414 | 160.0 | 14.38% | 5206.3 | 11.93% | motif file (matrix) | svg |
| 192 | C G T A G C T A C G T A G C A T A G C T A C T G A C G T C G T A A G T C A C T G A C T G G C T A G T C A C T G A G C T A | SPL14(SBP)/col-SPL14-DAP-Seq(GSE60143)/Homer | 1e-2 | -4.823e+00 | 0.0416 | 40.0 | 3.59% | 1041.0 | 2.39% | motif file (matrix) | svg |
| 193 | C G T A C G A T C G A T C G T A C T G A G T C A A G T C A G T C A C G T C G T A C T G A G C A T C G A T C G A T G C T A | AT1G72740(MYBrelated)/colamp-AT1G72740-DAP-Seq(GSE60143)/Homer | 1e-2 | -4.780e+00 | 0.0432 | 194.0 | 17.43% | 6453.2 | 14.79% | motif file (matrix) | svg |
| 194 | A C T G C T A G A G T C A C T G A C T G A T G C A C T G T A C G | ESE1(AP2EREBP)/col-ESE1-DAP-Seq(GSE60143)/Homer | 1e-2 | -4.778e+00 | 0.0432 | 284.0 | 25.52% | 9791.5 | 22.45% | motif file (matrix) | svg |
| 195 | A T C G A G T C A C T G A G T C A G T C A C T G G A C T G A C T | PUCHI(AP2EREBP)/colamp-PUCHI-DAP-Seq(GSE60143)/Homer | 1e-2 | -4.772e+00 | 0.0432 | 263.0 | 23.63% | 9008.0 | 20.65% | motif file (matrix) | svg |
| 196 | A C T G A C T G A G T C A C T G A C T G A G T C A C G T T C A G | ERF2(AP2EREBP)/colamp-ERF2-DAP-Seq(GSE60143)/Homer | 1e-2 | -4.749e+00 | 0.0439 | 265.0 | 23.81% | 9086.7 | 20.83% | motif file (matrix) | svg |
| 197 | G A C T C A G T G C A T C G A T T G A C A C G T A T G C G T A C C T G A A C T G A C T G A G C T | WIP5(C2H2)/colamp-WIP5-DAP-Seq(GSE60143)/Homer | 1e-2 | -4.745e+00 | 0.0439 | 259.0 | 23.27% | 8863.9 | 20.32% | motif file (matrix) | svg |
| 198 | A T G C A G T C G C T A C G A T C G T A G C A T G C T A C G A T C T A G C A T G T G A C G T C A | CArG(MADS)/PUER-Srf-ChIP-Seq(Sullivan\_et\_al.)/Homer | 1e-2 | -4.718e+00 | 0.0448 | 57.0 | 5.12% | 1605.1 | 3.68% | motif file (matrix) | svg |
| 199 | C T G A C G A T C A T G A T C G G C A T C A T G G C T A A G T C | ASHR1(ND)/col-ASHR1-DAP-Seq(GSE60143)/Homer | 1e-2 | -4.706e+00 | 0.0452 | 264.0 | 23.72% | 9057.1 | 20.76% | motif file (matrix) | svg |
| 200 | C G T A C T A G C A G T A C G T C G T A A C T G C A T G G C A T T C A G C T G A | MYB49(MYB)/col-MYB49-DAP-Seq(GSE60143)/Homer | 1e-2 | -4.704e+00 | 0.0452 | 179.0 | 16.08% | 5917.4 | 13.56% | motif file (matrix) | svg |
| 201 | T A C G T C A G G A T C G T A C T C G A G A C T G C T A G C T A G C T A C G T A G A T C G T C A | CDX4(Homeobox)/ZebrafishEmbryos-Cdx4.Myc-ChIP-Seq(GSE48254)/Homer | 1e-2 | -4.703e+00 | 0.0452 | 127.0 | 11.41% | 4039.3 | 9.26% | motif file (matrix) | svg |
| 202 | T C G A T A G C G T C A A C T G C T A G C G T A C G A T A C T G A C G T A C T G A C T G A C G T | ETS:RUNX(ETS,Runt)/Jurkat-RUNX1-ChIP-Seq(GSE17954)/Homer | 1e-2 | -4.679e+00 | 0.0457 | 32.0 | 2.88% | 796.2 | 1.83% | motif file (matrix) | svg |
